# Evolutionary design of regulatory control. I. A robust control theory analysis of tradeoffs

**DOI:** 10.1101/332999

**Authors:** Steven A. Frank

**Affiliations:** Department of Ecology and Evolutionary Biology, University of California, Irvine, CA 92697–2525, USA

**Keywords:** Phenotypic plasticity, homeostasis, systems biology, feedback

## Abstract

The evolutionary design of regulatory control balances various tradeoffs in performance. Fast reaction to environmental change tends to favor plastic responsiveness at the expense of greater sensitivity to perturbations that degrade homeostatic control. Greater homeostatic stability against unpredictable disturbances tends to reduce performance in tracking environmental change. This article applies the classic principles of engineering control theory to the evolutionary design of regulatory systems. The engineering theory clarifies the conceptual aspects of evolutionary tradeoffs and provides analytic methods for developing specific predictions. On the conceptual side, this article clarifies the meanings of *integral control, feedback*, and *design*, concepts that have been discussed in a confusing way within the biological literature. On the analytic side, this article presents extensive methods and examples to study error-correcting feedback, which is perhaps the single greatest principle of design in both human-engineered and naturally designed systems. The broad framework and associated software code provide a comprehensive how-to guide for making models that focus on functional aspects of regulatory control and for making comparative predictions about regulatory design in response to various kinds of environmental challenge. The second article in this series analyzes how alternative regulatory designs influence the relative levels of genetic variability, stochasticity of trait expression, and heritability of disease.

## Introduction

Regulatory control adjusts the expression of labile characters. The study of labile characters has developed along two separate lines. Molecular systems biology and physiology emphasize the biochemical mechanisms and immediate response of observable systems. Evolutionary biology analyzes how phenotypically labile or plastic characters influence variability in populations and ultimate reproductive function, causing change in the design of organisms.

Studies rarely combine the details of regulatory control architecture with the evolutionary analysis of variability and change in populations. In this article, I work toward building the theoretical foundation for integrating regulatory control and evolutionary perspectives (Koonin & Wolf, 2006; Soyer, 2012).

In the mechanistic literature, studies in systems biology, physiology, and behavior consider how regulatory control systems respond to changes in the environment. These disciplines have rich theories about adjustable phenotypes (Mazur, 2006; Alon, 2007a; Keener & Sneyd, 2009; Ingalls, 2013).

In the evolutionary literature, studies focus on the association between characters and reproductive fitness, the correlation between characters, the processes that influence genetic and phenotypic variation in populations, the evolutionary dynamics of genes and characters, and the consequences of lability or plasticity for the evolutionary origins of new characters and new species (Pigliucci, 2001; DeWitt & Scheiner, 2004; West-Eberhard, 2005).

Lande (2014) emphasized that most evolutionary theories of plasticity focus on how characters are set during a brief critical period of development. Few theoretical studies have analyzed the evolution of labile characters that adjust continuously throughout an organism’s lifetime (Mangel & Clark, 1988; McNamara & Houston, 1996; Frank, 2002; Fischer et al., 2014). How can we broaden the insights of evolutionary analysis to include the rich array of labile characters at the molecular, physiological, and behavioral level?

As a first step, I will use the general approach of engineering control theory to describe the universal features of regulatory control architecture (Iglesias & Ingalls, 2009). Control theory allows us to relate particular design aspects of regulatory control to evolutionary problems. For example, error-correcting feedback is a general design property common to many regulatory control systems. How can we relate error-correcting feedback to evolutionary aspects of design tradeoffs between different components of performance? What are the consequences of those tradeoffs for genetic variability and stochasticity in phenotypic expression?

In this series of articles, I combine the methods and insights of engineering control theory with the evolutionary analysis of labile characters. This first article introduces the methods and analyzes fundamental design tradeoffs for labile characters. The second article in this series studies the consequences of alternative control architectures for genetic variability, phenotypic stochasticity in trait expression, and the heritability of disease (Frank, 2018b). Further articles develop the interplay between control architecture and evolutionary dynamics.

## Overview

This article analyzes evolutionary design tradeoffs for regulatory control systems. To develop basic concepts, I focus on two design goals. First, how do organisms track changing environmental signals with plastic, responsive regulatory control? Second, how do organisms maintain a homeostatic setpoint in spite of environmental perturbations?

Those two goals typically trade off, because faster responsiveness often improves environmental tracking but degrades homeostatic maintenance. The tradeoff between plastic responsiveness and homeostatic maintenance depends on additional performance tradeoffs, which I develop throughout this article.

I emphasize the analytic methods by which one can model the various tradeoffs and make predictions about organismal design. I also discuss how one should think about the fundamental concepts of regulatory design.

Initially, I focus on functional design aspects of control, such as error-correcting feedback, rather than on mechanistic aspects, such as how particular molecules change in expression. Ultimately, the theory must merge functional and mechanistic perspectives in particular applications. However, an evolutionary foundation must begin with the basic framing of function.

On the functional side, error-correcting feedback is perhaps the single greatest design principle in both human-engineered and naturally evolved control systems. Yet, the biological literature on error-correcting feedback control presents various and sometimes conflicting meanings of *feedback*. To clarify the meaning of *feedback*, one has to have a clear notion of the meaning of *design* in biological systems.

By developing those broad analytic and conceptual topics, this article presents a how-to guide for making and interpreting evolutionary models of regulatory control in biology.

The first section begins with a simple model for tracking a changing environmental signal. The second section introduces the methods of engineering control theory. That theory provides the most powerful tools for analyzing and interpreting regulatory control with respect to design goals, such as tracking and homeostasis.

The third section presents some alternative mechanisms by which a biological system could control a process to achieve a design goal. One typically begins with some intrinsic dynamical process, such as a biochemical reaction, and then considers how an organism modulates those given dynamics to improve performance. Control designs include integrating deviations from target dynamics, feeding back error into the system so it can self-correct, and using filters to reject unwanted inputs.

The fourth section clarifies key concepts of control that are often discussed in a confused way within the biological literature. Those concepts include *integral control, feedback*, and *design*.

The fifth and sixth sections summarize major design tradeoffs and present performance measures that can be used to model those tradeoffs. I emphasize the tradeoffs among the plastic responsiveness of environmental tracking, the homeostatic rejection of perturbations, system stability, and the costs of controls that modulate dynamics.

I also consider how the various performance goals may trade off with robustness, which is the reduced sensitivity to random perturbations and other uncertainties. Another key tradeoff concerns performance in relation to different frequencies of inputs. Better performance to slowly changing inputs may trade off against poorer performance with respect to more rapidly changing inputs.

The seventh section analyzes the dynamics of an intrinsic biological process, such as a biochemical reaction. Study of an intrinsic process in the absence of modulating control sets the stage for the eighth section, which provides a detailed analysis of how error-correcting feedback control can modulate the intrinsic process and improve performance.

The analysis of performance leads to explicit models of the various design goals and tradeoffs of control systems. I develop optimization methods and present several analytical and numerical examples.

The ninth through twelfth sections emphasize particular design tradeoffs. Each section presents analytic methods and numerical examples. The tradeoffs with performance include the costs of control, the balance between plastic responsiveness and homeostatic maintenance, the costs and benefits of error feedback versus simpler direct control architectures, and the consequences of improving system stability subject to the cost of reduced performance.

The final section discusses extensions and conclusions. The supplemental files provide all of the software code in Wolfram Mathematica and C++ used to develop the analysis, numerical examples, and graphics. That code includes details of the methods and analysis. The code also forms the basis for developing novel research projects on regulatory control.

## Environmental signal tracking

Lande’s (2014) simple exponential model for environmental tracking provides a good way to link the current evolutionary literature of phenotypic plasticity to the concepts and methods of engineering control theory.

Following Lande, suppose that the labile component of phenotype tracks the environment by the simple differential equation

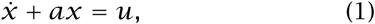

in which the overdot denotes differentiation with respect to time, *t*, the term *x*(*t*) is the phenotypic deviation from the initial condition, *x*(0) = 0, and phenotypic deviations are driven by the environmental input signal, *u*(*t*). I use notation that provides a natural connection to control theory. The relation to Lande’s notation is *x*(*t*) ≡ *ξ_t_, a* ≡ *λ*, and *u*(*t*) ≡ *λϕ*(*∊_t_*).

By standard analysis, the phenotypic deviation from the initial baseline at time *t* is

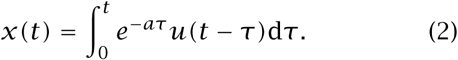

This process describes how the phenotypic deviation, *x*, arises from the sequence of environmental input signals, *u*(*τ*), and the intrinsic rate of decay for phenotypic deviations, *a*.

If we know the dynamics of the environmental signal, *u*(*τ*), we can use the integral solution to calculate *x*(*t*). However, such calculations can be tedious and often provide little insight. For example, we might ask, in general, how an increase in the rate of tracking adjustment, *a*, influences the benefit of closely tracking the environment versus the cost of responding too strongly to noisy signals or over-adjusting to sudden environmental shifts.

## Control theory analysis

The classic control theory approach to the analysis of dynamics provides much easier calculations of system response and much deeper insight into general aspects of tradeoffs in the design of responsive systems. In addition, control theory analysis encourages a more explicit description for the mechanistic basis of biological regulatory systems. With an explicit description of regulatory control, one can connect the specific design tradeoffs for phenotypes to the underlying consequences for genetic variability and the stochastic aspects of phenotypic expression.

This section briefly reviews two key aspects of classic control theory. I follow my recent tutorial exposition (Frank, 2018a). Additional details can be found in standard texts of control theory (e.g., Åström & Murray, 2008; Ogata, 2009; Dorf & Bishop, 2016). The following sections apply these control theory analytical methods to evolutionary aspects of phenotypic plasticity, emphasizing the key tradeoffs that influence design. Throughout this article, the references in this paragraph can be used to follow up on technical aspects of signal processing and control theory.

### Transfer functions

Transfer functions are a particularly important tool in control theory analysis. Here, I list some of the basic notation and how one can use transfer functions to analyze dynamics. Good introductions can be found in most textbooks on control theory (e.g., Åström & Murray, 2008; Ogata, 2009; Dorf & Bishop, 2016). My tutorial provides many basic examples and discusses limitations and potential remedies (Frank, 2018a).

We can transform the temporal dynamics of any linear time-invariant differential equation in the time variable *t* into an expression in the complex Laplace variable *s*. For example, given the differential equation in the time variable *t* as

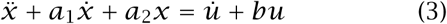

with *y ≡ x* (see below), we can write the dynamics equivalently with functions of the complex Laplace variable *s* as

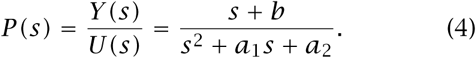

The numerator expresses a polynomial in *s* derived from the coefficients of *u* in eqn 3. Similarly, the denominator expresses a polynomial in *s* derived from the coefficients in *x* from the left side of eqn 3. The eigenvalues for the process, *P,* are the roots of *s* for the polynomial in the denominator.

From eqn 4 and the matching picture in Fig. 1a, we may write *Y*(*s*) = *U*(*s*)*P*(*s*). In words, the output signal, *Y*(*s*), is the input signal, *U*(*s*), multiplied by the transformation of the input signal by the process, *P*(*s*). Because *P*(*s*) multiplies the signal, we may think of *P*(*s*) as the signal gain or amplification, which is the ratio of output to input, *Y*/*U*.

**Figure 1:**
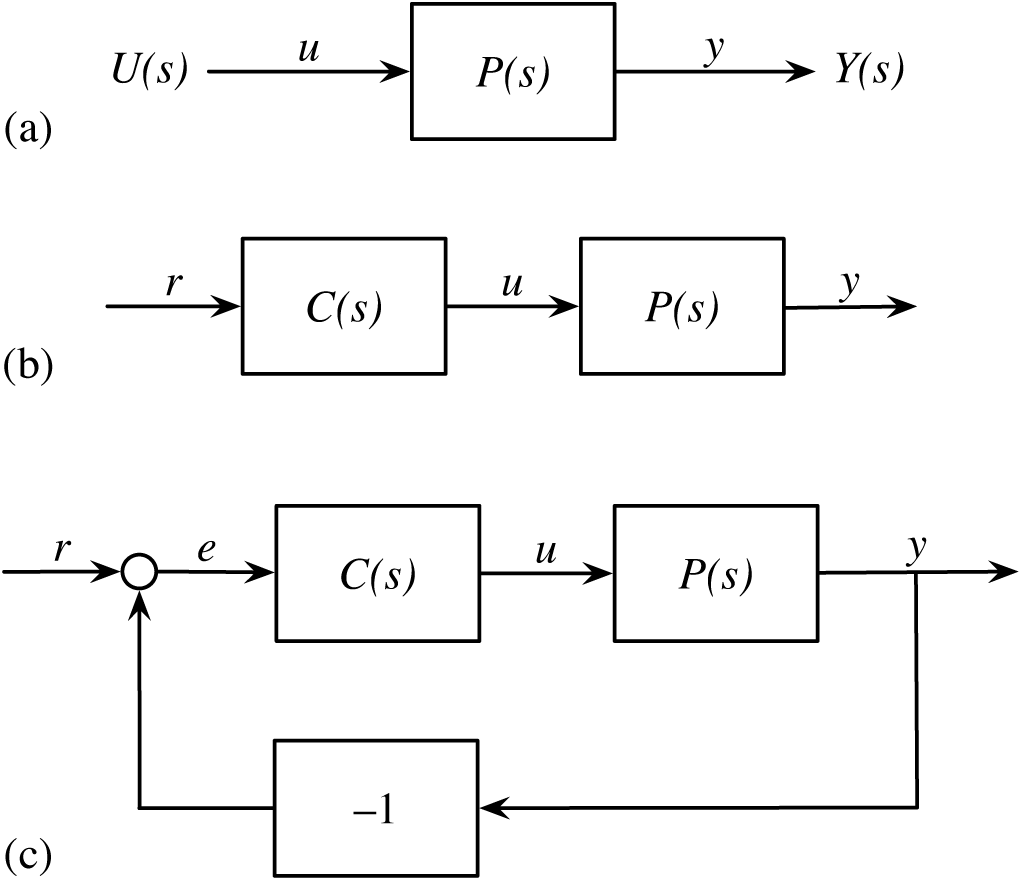
Mechanistic descriptions of control. (a) The input-output flow in eqn 4. The input, *U*(*s*), is itself a transfer function. However, for convenience in diagramming, lower case letters are typically used along pathways to denote inputs and outputs. For example, in (a), *u* can be used in place of *U*(*s*). In (b), only lower case letters are used for inputs and outputs. Panel (b) illustrates the input-output flow of eqn 5. These diagrams represent open loop pathways, because there is no closed loop feedback pathway that sends a downstream output back as an input to an earlier step. (c) A basic closed loop process and control flow with negative feedback. The circle between *r* and *e* denotes addition of the inputs to produce the output. In this figure, *e = r − y*, is the error between the environmental reference input, *r,* and the system output, *y*. From Frank (2018a).

Following the conventions of control theory, the system output *Y* expresses a transformation of the internal system state, *X*. In our initial examples, the output is equivalent to the internal system state, *y*(*t*) ≡ *x*(*t*), and thus *Y*(*s*) ≡ *X*(*s*).

The simple multiplication of the signal by a process means that we can easily cascade multiple input-output processes. For example, Fig. 1b shows a system with extended input processing. The cascade begins with an initial reference input, *r,* which is transformed into the command input, *u*, by a preprocessing controller, *C*, and then finally into the output, *y*, by the intrinsic process, *P.* The input-output calculation for the entire cascade follows easily by noting that *C*(*s*) = *U*(*s*)/*R*(*s*), yielding

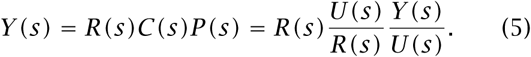

These functions of *s* are called *transfer functions*.

The transfer function in eqn 4 includes the exponential tracking model in eqn 1 as a special case. We can write that transfer function for eqn 1 as

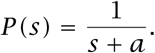

We can always multiply *P* by any constant to change the output by that constant value. So we may choose to multiply *P* by *a* and write the exponential tracking model as

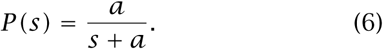

This modified form has the benefit that as *s* → 0, the gain of the process, *P,* goes to a normalized value of one. This normalized expression of *P* is the classic form of the basic low-pass filter, as described in the next section.

### Frequency domain

In a standard temporal analysis, we might begin with a description of dynamics, such as eqn 1, and ask how the system state *x*(*t*) changes with various fluctuating inputs, *u*(*t*). In control theory language, how do fluctuations in the input signal *u* influence the output signal, *x*?

A linear time-invariant system transforms a sine wave input into a sine wave output at the same frequency, but with altered magnitude and phase. Consider the response of the system in eqn 1, with associated transfer function eqn 6, to sine wave inputs of frequency, *ω*. The left column of panels in Fig. 2 illustrates the fluctuating output in response to the green sine wave input. The blue (slow) and gold (fast) responses correspond to parameter values in eqn 6 of *a =* 1 and *a =* 10.

**Figure 2:**
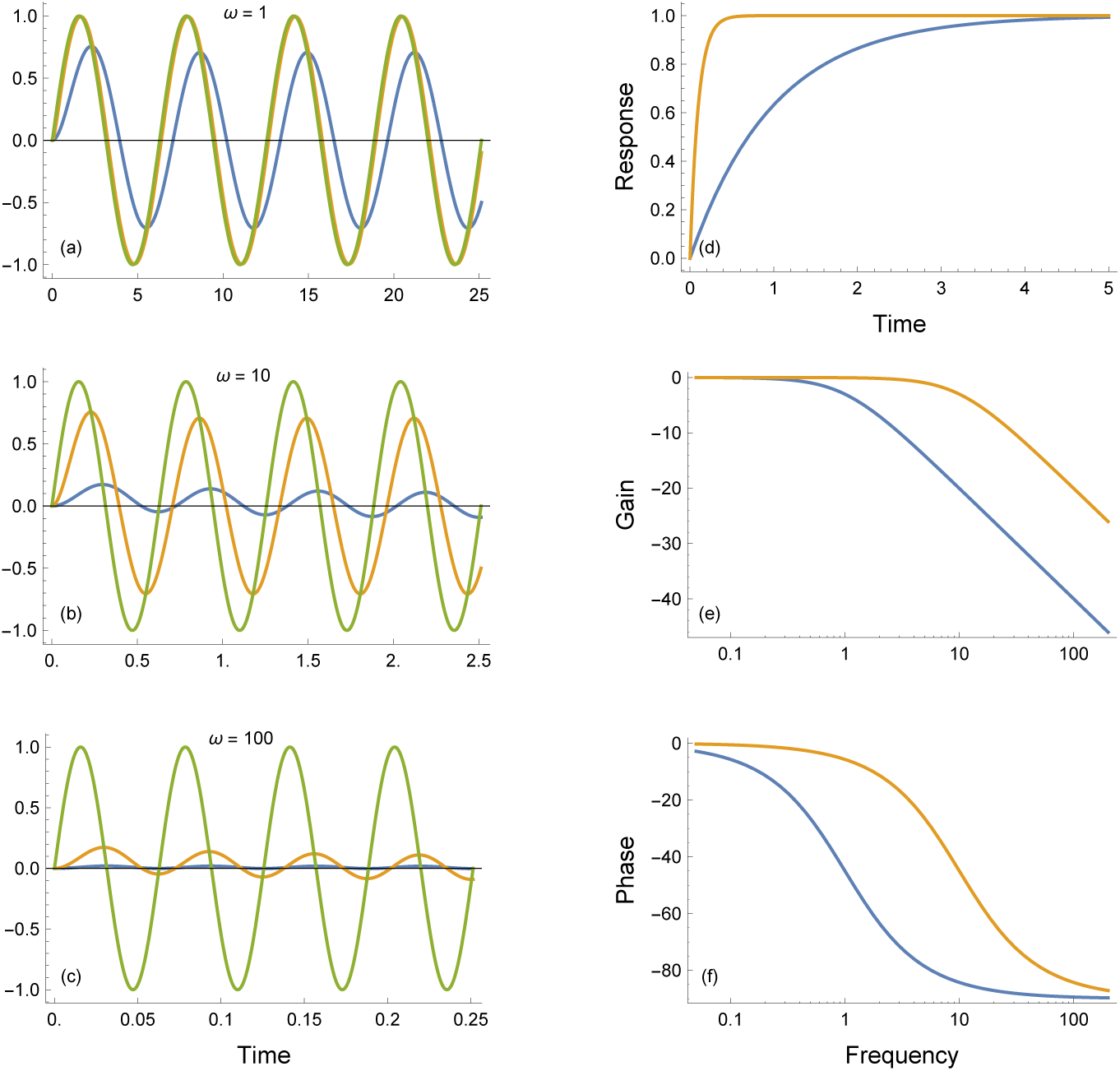
Response of the low pass filter in eqn 6. The blue (slow) and gold (fast) responses correspond to parameter values in eqn 6 of *a =* 1 and *a =* 10. (a-c) Temporal dynamics in response to input *u*(*t*) (green curve) as sine waves with varying frequencies, *ω*. (d) Response of eqn 6 to unit step input, *u*(*t*) = 0 for *t* < 0 and *u*(*t*) = 1 for *t* ≥ 0. (e) The output-input gain ratio for the transfer function in eqn 6 as a function of input frequency. This Bode plot shows the gain on a scale of 20 log_10_(gain). A log gain value of zero corresponds to a gain of one, log(1*) =* 0, which means that the output magnitude equals the input magnitude. (f) The phase shift of the output vs input sine waves as function of the input frequency, *ω*. This Bode phase plot shows the angular phase shift in degrees. Original figure and additional details in Frank (2018a).

Both the slow and fast transfer functions pass low frequency inputs into nearly unchanged outputs. At higher frequencies, they filter the inputs to produce greatly reduced magnitude, phase-shifted outputs. The transfer function form of eqn 6 is therefore called a low pass filter, passing low frequencies and blocking high frequencies. The two filters in this example differ in the frequencies at which they switch from passing low frequency inputs to blocking high frequency inputs.

The Bode gain plot in Fig. 2e provides a particularly important summary of a dynamical system’s response to fluctuating inputs. The gain is the ratio of the output magnitude to the input magnitude, the amount by which the transfer function amplifies its input. The Bode plot shows a transfer function’s gain at various input frequencies.

## Mechanisms of phenotypic response

The simple exponential tracking model in eqn 1 can be related to different underlying mechanisms that control phenotypic response to the environment. This section describes three alternative mechanistic systems of control. The alternative mechanisms have different evolutionary consequences. The alternatives also highlight fundamental principles that apply broadly to regulatory control systems in biology. The following sections analyze these three different mechanistic interpretations.

### Uncontrolled process

The exponential response to the environment may be an intrinsic aspect of the organism. Figure 3 illustrates an intrinsic exponential process, *P.* In control theory, *P* typically signifies an unmodifiable “plant” process.

**Figure 3:**
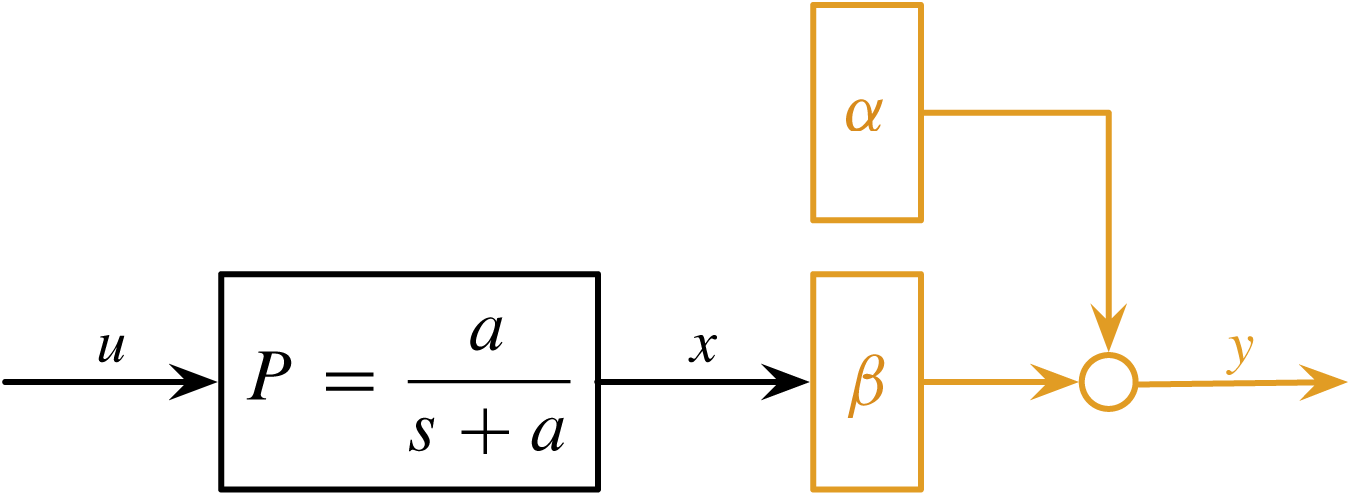
An exponential process response, *P,* with the output signal modified by postprocessing. The black components are intrinsic aspects that cannot be modified. The gold components are the affine transformation of the intrinsic process output, *x*, to yield the final output, *y = α + βx*. The variables *α* and *β* of the affine transformation can be genetically variable and modified by evolutionary processes. This description of an exponential system with genetically variable affine postprocessing matches Lande’s (2014) model.

Within the scope of our biological analysis, we consider the intrinsic process to be constrained and not subject directly to change. Any modification of the organism’s response must arise by preprocessing the input signal that comes into *P* or postprocessing the output signal produced by *P.*

Figure 3 illustrates Lande’s (2014) model, in which the black components show the environmental input, *u*, and the intrinsic exponential organismal response, *x*. The gold postprocessing yields the final response, *y = α + βx*, in which the postprocessing can be modified by natural selection through the genetically variable traits *α* and *β*.

In this case, the component of phenotypic lability subject to natural selection is the affine transformation of the intrinsic response *x*, into the final organismal response, *y*. *Affine transformation* simply means a constant shift and stretch, here a shift by *α* and a stretch (or shrink) by *β*.

An intrinsic exponential response may arise by a relatively simple process. If the rate of increase in the response depends on the input, 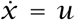, and the response degrades at a rate proportional to the current response level, 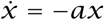, then we obtain the basic differential equation for the exponential response 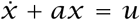, as given in eqn 1. For example, *x* may be a molecule produced at a rate influenced by an incoming signal, *u*, and degraded at a constant rate, *a*. Such stimulus-triggered production and intrinsic degradation describe the most basic biochemical system response.

### Integral control and feedback

Suppose an organism’s intrinsic “plant” response to input simply mimics the input level. For example, organismal surface temperature may closely track the ambient temperature. Figure 4a illustrates an intrinsic “plant” with *P =* 1, a simple pass-through process in which the output *y = uP* is equal to the input *u*.

**Figure 4:**
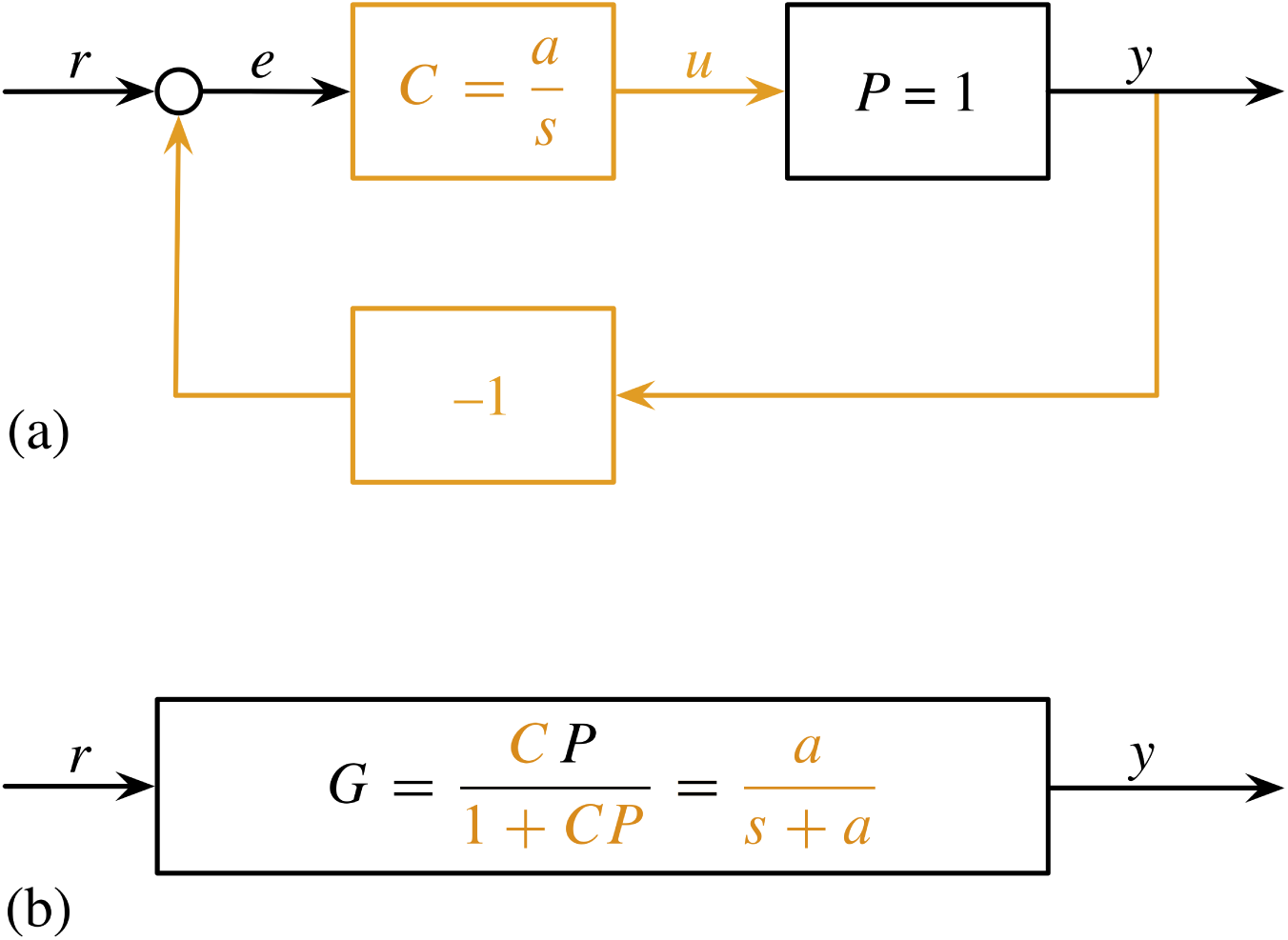
Feedback loop with an integral controller. (a) The black box is fixed as an intrinsic process, the gold components can be adjusted by evolutionary or other design processes. (b) The entire feedback loop can be collapsed into a single transfer function and associated box, *G*, which is the exponential process. The denominator of *G* represents the feedback loop component, which is a designed feature, and so is entirely in gold.

Such instantaneous tracking of the environment has the benefit of quick adjustment to external change. But rapid adjustment can also be costly. Short-term rapid external fluctuations may simply be noise in the input, such as fluctuations in light intensity that cause rapid shifts in the local surface temperature. Organisms often benefit by ignoring very rapid, noisy fluctuations, and tracking slower changing, more reliable signals of the external environment.

In addition, the intrinsic pass-through process may have stochastic error, *δ*, with an associated plant, *P =* 1 *+ δ*. Ideally, the organism could correct for such intrinsic fluctuations and potential unknown internal biases.

How can an organism track the slower, more reliable external signals, reject the noisy external fluctuations, and adjust to any biases or fluctuations in the internal pass-through plant process?

Figure 4a shows how an organism can modulate its response through two genetically modifiable components, shown in gold.

First, the signal coming into the intrinsic plant may be altered by a controller, *C*. In control theory, the controller can be modified by design to alter the input signal, *u*, passed into the intrinsic plant process. Here, we assume “design” means evolutionary processes, such as natural selection, subject to the constraint that the controller can only take on forms that can be realized by the organism’s physiology and genetics.

Second, the organism’s final output response, *y = x*, is fed back into the system as an additional input. By feeding back the output and subtracting that feedback from the external environmental signal, now labeled as *r* for the external reference, the actual value that enters the first preprocessing controller step is *e = r* − *y*. That value is the error difference between the external environmental reference signal and the actual output by the system.

Error feedback is perhaps the single most powerful mechanism of system design in both human-engineered and naturally evolved systems. By feeding the error into the system as its primary input, the system can always correct any perturbations and misspecifications by moving to reduce the error. If the error is positive, the system moves to increase the output. If the error is negative, the system moves to reduce the output. With error-correcting feedback, a sloppy, poorly specified system can still perform well.

Now consider the first modifiable component, the controller. To transform the error input, *e*, into the control command, *u*, the organism could potentially use any process that can be realized physiologically and genetically. Here, in order to develop the alternative interpretations of the fundamental exponential model, I confine the controller to be the transfer function, *C*(*s*) = *a*/*s*, with modifiable parameter, *a*. The transfer function 1*/s* is a pure integrator, because it corresponds to the differential equation 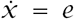 for input *e* and internal state, *x*. Thus, the value of the internal state is the integral of the error input, 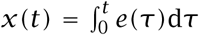. The *a* term in the numerator of *C = a/s* multiplies the integral output of *C* by the constant value, *a*.

Another compelling benefit of transfer functions is that we can easily calculate the total system response of a feedback loop. If we write all signals and internal processes as transfer functions, with *Y* as the transfer function for the output signal, *y*, and *E* as the transfer function for the error signal, *e*, then the direct line of signal processing between the input and the output without feedback yields an output *Y = CPE*, because transfer functions multiply along a signal line. Noting that *E = R − Y* and substituting that expression into the previous input-output expression, we obtain

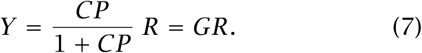

The complete feedback loop system, *G*, that takes input *R* and yields output *Y* is

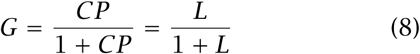

in which *L = CP* is often called the open loop component of the system—the open part of the system without the feedback that closes the loop. In this case, we have *C = a/s* and *P =* 1, thus *G* = *a*/(*s* + *a*) is our basic exponential process in eqn 6. In general, the transfer functions *C* and *P* can describe any linear time-invariant system.

### Low-pass preprocessing filter

An organism may perceive the environment through a sensor, which transforms the environmental input signal. In Fig. 5, the sensor or preprocessing filter, *F,* transforms the input, *r,* by our standard exponential process. The filtered signal, *f,* then enters the organismal system, where it may be further processed by the system, *G*. I show *G* as a feedback loop in this example.

**Figure 5:**
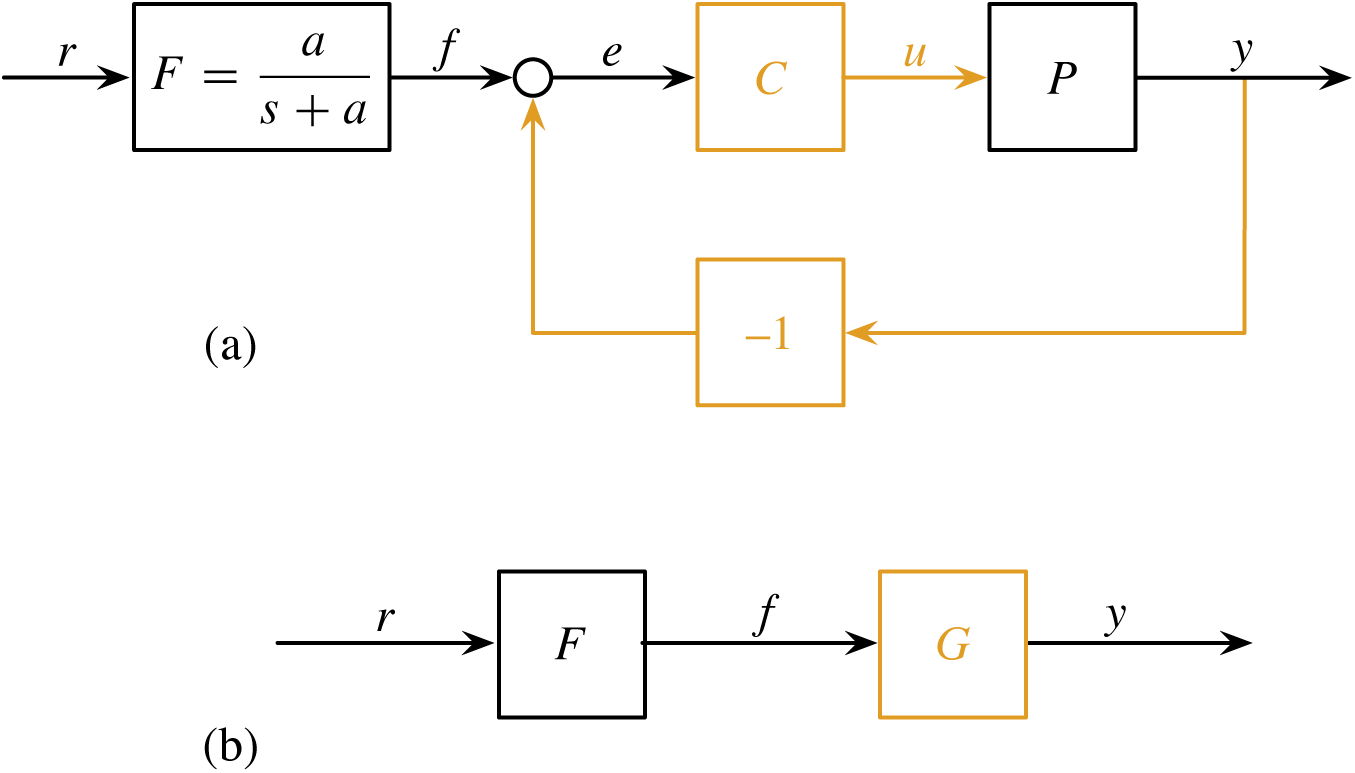
The exponential process as a preprocessor, *F,* that filters the environmental reference signal before entering a postprocessing feedback loop. (a) Here, *F* is assumed to be a fixed aspect of the organism, such as an unmodifiable sensor of the environment. In other cases, we may consider *F* as a modifiable designed filter of system input. The postprocessing feedback loop includes an unmodifiable intrinsic plant process, *P,* and a modifiable controller and feedback process. (b) We can collapse the post-processing feedback loop into a single transfer function and associated box, *G*. We may interpret *G* as a non-feedback description of a dynamical process or as a feedback loop. Our interpretation depends on whether we consider the tendency to attract toward the reference input as an intrinsic unmodifiable aspect of the dynamics or as a modifiable feedback feature designed to move the system toward the reference value at a particular rate.

The feedback loop in Fig. 5a contains the standard components: a intrinsic process, *P,* that cannot be modified, a modifiable controller, *C*, and an optional feedback loop that is included among the modifiable components of the system. This postprocessing system may include an integral component in the controller of the feedback loop.

## Interpretation of integral control and feedback

Integral control and feedback are key concepts in control theory. Those key concepts are sometimes misunderstood when analyzing systems designed by natural biological processes. Consider, for example, a system in which the production rate of some entity is proportional to the input, and the production rate is balanced by a matching degradation rate. The dynamics follow the simple exponential process of eqn 1, with transfer function in eqn 6.

Should we interpret that exponential process as a designed system with integral control and error-correcting feedback or as a simple unitary component with an exponential response? That broad question raises three specific questions.

### What is integral control?

An intuitive understanding of integral control can be obtained from eqn 7 and 8. If the input reference signal, *R*, is a constant, then the system can match its output to the input and reduce the error to zero only if *G =* 1. For *G* → 1, we must have *L* → *∞*. In practice, we must have a very high amplification of the input signal by the open loop, *L*.

The higher the open loop gain, the more strongly a feedback system drives the error toward zero. We can describe that role of high open loop gain in a feedback system directly by expressing the error, *E = R* − *Y,* from eqn 7 and 8 as

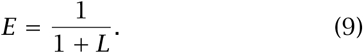

This equation expresses one of the great principles of design. High open-loop amplification of a signal drives the error to zero in a feedback loop. Powerful error-correcting feedback compensates for many kinds of perturbations, uncertainties, and sloppy components that perform poorly. Nonlinearities can often be thought of as uncertainties in linear system dynamics. Thus, an error-correcting feedback system designed as if it were linear often performs well if the actual dynamics follow particular kinds of nonlinearity (Vinnicombe, 2001; Frank, 2018a).

How do we obtain a very high gain for the open loop, *L*, when the input signal is constant? If *L* has an integrator component, 1*/s*, then *s* → 0 implies *L* → *∞*. A temporally constant reference input associates with a zero frequency input, which corresponds to *s =* 0 in a standard analysis of sine wave inputs. Thus, *L* must have an integral component in order for the system to achieve a perfect match to a constant input signal.

The recent systems biology literature on responsive biochemical processes has elevated integral control to an almost mythical status by which biological systems achieve a perfect matching response to transient environmental inputs, often called “perfect adaptation” (Yi et al., 2000). Although true, one must understand three key points.

First, feedback is a powerful error-correcting design feature that typically requires high gain.

Second, to achieve high gain at low input frequency, *L* must increase for small *s* values, which correspond to low frequency inputs. A pure integrator 1*/s* goes to infinity at zero frequency. In practice, high gain at low frequency is sufficient.

Third, integral control simply means that, for some component of the system, the production rate of a molecule or other physical entity is proportional to the input signal. That production is typically balanced by a degradation process. Proportional production and balancing degradation arise often in biochemical systems and form the most basic type of feedback loop, creating exponential components. Thus, integral control and “perfect adaptation” are not mystical achievements, but instead are common outcomes of basic feedback dynamics.

### What is feedback?

A process that balances production and degradation may be thought of as a feedback system. However, a process that produces heat and passively dissipates that heat would not typically be thought of as a feedback process to regulate heat. When should we consider balancing forces as feedback (Fig. 4a), and when should we consider the same dynamics as a unitary component of system dynamics (Fig. 4b)?

Recent systems biology analyses of phenotypically responsive biochemical interactions add further confusion to the meaning of *feedback.* For example, the classification of molecular network motifs and control architecture sometimes use *feedback* to describe a system in which the concentration of one molecule influences the dynamics of a second molecule, and the concentration of the second molecule in turn influences the dynamics of the first molecule. That notion of feedback emphasizes the mutual influence between physical entities (Alon, 2007a,b).

Control theory emphasizes an alternative, abstract notion of feedback, in which system dynamics can be interpreted as in Fig. 4a. In that abstract interpretation, there is some reasonable way in which to describe the system as subtracting the output from the external input and then feeding that error difference as the input into the system.

The confusion in the systems biology literature is particularly strong, because that subject emphasizes both the abstract control theory analysis of systems and the incompatible interpretation of feedback as mutual influence between physical entities (Alon, 2007a). Until one realizes the distinction between the alternative interpretations of feedback, the literature can be difficult to understand. However, it is worthwhile to sort things out, because systems biology has developed the most comprehensive conceptual and mechanistic analyses of phenotypically responsive systems. A clear notion of *design* helps.

### What is a designed system?

When should we describe a balance between production and degradation by the special design concepts of integral control and feedback?

In general, a modifiable component that has been tuned to achieve some goal forms part of a designed system. The goal may be a target that has been set within a human-engineering context. Or the goal may be the consequence of natural biological processes that shape phenotypes as if they were designed to achieve particular functions favored by natural selection.

George Williams (1966) clarified the biological interpretation of design. In Williams’ view, if a fish jumps out of the water and then returns to the water by gravity, we would not consider the path of return a designed feature. Gravity is a sufficient explanation. By contrast, if the fish uses its modified fins as sails to slow the rate of return, then the use of fins as sails is a designed feature to achieve the goal of altering the path of return to the water.

What about a simple biochemical system of production and degradation that acts as an exponential process? If the production rate has been modified to achieve high amplification in response to low frequency input signals, then that integral control aspect may be thought of as a designed feature. By contrast, an intrinsic reaction that passively changes its production in response to changing ambient temperature may be thought of as an inevitable physical consequence of the input energy.

With regard to feedback, degradation rates can be modified by reactions that specifically destroy a molecule or by changes in molecular structure that alter the rate of degradation. If the degradation rate has been modified to balance production and track some target setpoint of molecular abundance, then degradation acts as a designed feedback mechanism. By contrast, an intrinsic decay rate may be thought of as an inevitable physical process rather than as a feedback design.

## Design tradeoffs

I focus on the tradeoffs faced by evolutionary processes in the design of organisms. This section lists some of the key biological tradeoffs, expressed in terms of the methods and insights of engineering control theory.

Designed systems typically have multiple goals. For example, an organism gains by adjusting to environmental changes. It also gains by homeostatically holding its internal state in response to noisy environmental fluctuations.

Often, there may be tradeoffs between alternative goals. The more rapidly an organism changes its state to track environmental changes, the more susceptible it may be to environmental fluctuations that disrupt the organism’s internal homeostasis. In other words, there may be a tradeoff between the phenotypic plasticity of environmental tracking and homeostatic regulation.

Other tradeoffs arise. The rate of system adjustment to external environmental changes may influence the tendency of the system to become unstable. Instability of a critical system may lead to death. The costs of building and running additional control structures must be balanced against the added benefits provided by those extra controls.

To summarize, four common design goals often tradeoff against each other: environmental tracking, homeostatic regulation, stability, and the costs of control. We may add robustness as a fifth design goal, in which robustness means reduced sensitivity to random perturbations and other uncertainties.

## Measures of performance

To analyze design with respect to tradeoffs, we must have measures of performance. In biology, one usually uses various measures of reproductive success or fitness as the ultimate measure of performance. Analysis of phenotypic plasticity requires explicit assumptions about how tracking, regulation, stability, and costs translate into the ultimate measure of reproductive performance.

I use the classic quadratic measure of performance from control theory (Anderson & Moore, 1989). Tracking considers how far the actual system output, or phenotype, is from the optimum. The usual performance measure sums up the Euclidean distances between the optimum and the actual output at each point in time as the squared errors, *e* = (*r − y*)^2^, between the target reference signal, *r*(*t*), and the system output, *y*(*t*). Summing over all infinitesimal time intervals from an initial time *t =* 0 to a final time *T* yields the quadratic measure

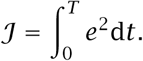

Optimal performance minimizes the total error, *𝒥*, subject to any processes that constrain the system.

Homeostatic regulation concerns deviations from a constant target value. We can, without loss of generality, take the target to be *r*(*t*) ≡ 0. Thus, the performance metric for homeostatic regulation is *𝒥*, in which *e*^2^ becomes *y*^2^, the squared deviation of the phenotype from the constant setpoint.

Tradeoffs often arise between tracking and homeostatic regulation. The more rapidly a system can track changes in the target reference signal, *r*(*t*), the more strongly a system tends to deviate from a constant homeostatic setpoint in response to random perturbations. Put another way, a beneficial response to a true change in the environmental input signal can also lead to a detrimental response to a false, noisy fluctuation in environmental or other inputs.

Typically, we also wish to consider the costs of control. For example, driving a system toward its optimum value often requires energy and other investments. The greater the energy expended per unit time, the faster the system can drive its output toward its optimum. However, performance must also consider minimizing the costs of energy expended and other investments in control. Thus, the classic performance measure of control is typically written as

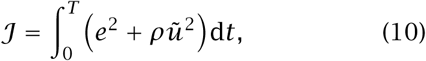

in which ũ(*t*) is a function of the magnitude of the signal, *u*(*t*), that the system uses to control its dynamics, as in Fig. 1c, and ∫*ũ*^2^d*t* is proportional to the total signal power. I use *ũ = u–r,* the difference between the control signal, *u*, and the input reference signal, *r.* This choice sets the cost for control to zero when the controller simply outputs the unchanged reference signal.

The parameter *ρ* weights the relative importance of the tracking error and the cost of control. This performance metric, *𝒥*, balances the tradeoff between minimizing the tracking error and minimizing the cost of control.

Stability sets an additional dimension of performance. For example, suppose we minimize *𝒥* to obtain an optimally controlled system with respect to the tradeoff between tracking error and control costs. Our optimal system may be prone to instability in response to small perturbations. Instability often leads to complete system failure.

To obtain a better control design, we often must impose a stability constraint on the minimization of performance, *𝒥*. For example, we may require that the system remain stable to particular kinds and magnitudes of perturbations. Such a constraint is often called a stability margin of safety, or simply a stability margin. Because instability is often disastrous, robust engineering design methods and natural biological design processes tend optimize performance subject to constraints on the stability margin.

Specifying a stability margin is technically more challenging than writing a simple performance metric (Vinnicombe, 2001; Frank, 2018a). I provide some examples in later sections.

We may also consider the robustness of various performance measures in relation to particular kinds of uncertainty. Greater robustness reduces the sensitivity of performance to uncertainty. However, reduced sensitivity may come with the cost of reduced maximum performance with respect to the target environment for which the design is tuned.

## Intrinsic plant process

I will illustrate the main design tradeoffs by a variety of examples. Each example typically begins with an intrinsic plant process that describes some fixed biochemical reaction or organismal input-output response. Given an intrinsic process, we may then consider how natural biological processes design regulatory control systems to modulate the intrinsic dynamics.

Before turning to the design problems, it is useful to have a clear sense of a simple set of generic intrinsic processes that can be used to build various initial examples. I focus on a basic second-order differential equation that is a reduced form of eqn 3, as

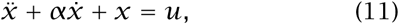

with associated transfer function

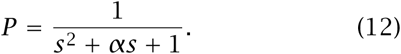

For *α ≥* 2, we can factor the denominator so that

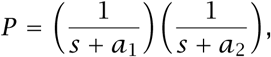

in which *a*_1_ *+ a*_2_ *= α* and *a*_1_*a*_2_ *=* 1. The factored expression shows that we can think of this system as the pair of cascading exponential processes shown Fig. 6a, because a cascade is described by the product of the transfer functions for each component. Algebraically, we can express the cascade by rewriting the second-order differential equation as a pair of firstorder equations

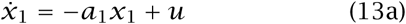

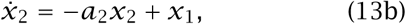

with system output *y = x*_2_. A cascade of exponential processes must be very common, because an exponential response arises from the most basic chemical process of production balanced by degradation.

**Figure 6:**
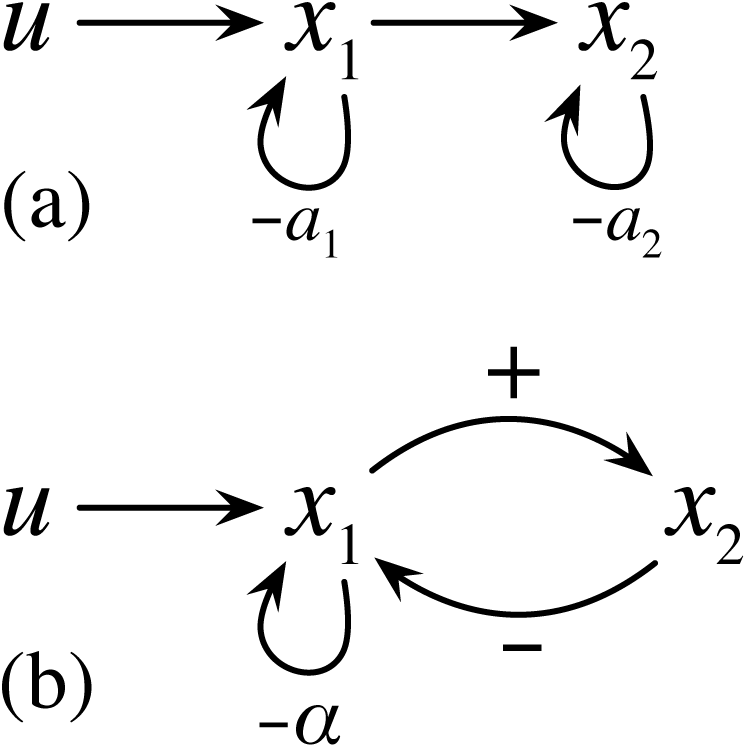
Examples of second-order dynamics mechanisms. (a) A cascade of exponential processes. The incoming signal *u* stimulates production of *x*_1_, which degrades at rate *a*_1_. The level of *x*_1_ stimulates production of *x*_2_, which degrades at rate *a*_2_. Dynamics given in eqn 13. (b) The first part of this mechanism is the same as the upper panel, with *u* stimulating production of *x*_1_, which degrades at rate *α*. In addition, *x*_1_ and *x*_2_ are coupled in a negative feedback loop. Dynamics given in eqn 14.

Alternatively, for any real value of *α*, including *α ≥* 2, we can rewrite the second-order differential equation as a pair of first-order equations

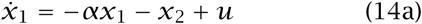

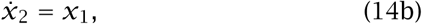

with system output *y = x*_2_. These first-order equations yield a simple graphical representation of a process that follows the dynamics of the secondorder system, as shown in Fig. 6b. The figure shows that this system can be described by a process with a negative feedback loop between two components, *x*_1_ and *x*_2_. Note that “feedback” in this sense describes a mechanistic interaction between two physical entities rather than the logical notion of “feedback” in a designed error-correcting control system.

When *α = u =* 0, the system is a pure oscillator that follows a sine wave. For 0 *< α <* 2 and *u =* 0, the system follows damped oscillations toward the equilibrium at zero, because the degradation of *x*_1_ at rate *–α* causes a steady decline in the amplitude of the oscillations about the equilibrium.

## Costs and benefits of error feedback

The examples in the prior section for the dynamics of *x*_1_ and *x*_2_ represent the intrinsic plant process, *P.* Figure 6 shows *u* → *P,* the control signal *u* as the input to the intrinsic process, *P.*

How can a system alter the dynamics of *P* in order to improve performance with respect to particular design goals? This section compares two basic approaches.

First, the system can modulate the control input signal, *u*, by modifying the dynamics of a controller process, *C*, as in Fig. 1b. In that system, the external environmental reference signal, *r,* enters as the system input into the controller, *C*, which outputs the control signal *u* that becomes the input into the intrinsic system process, *P,* which produces the final system output, *y*. That pathway is an open loop, because the system output, *y*, is not fed back as input to close the loop.

In the second approach to altering system dynamics, the output *y* is subtracted from the reference signal, *r,* and the resulting error *e = r* – *y* is fed back into the system as its input. The signal processing pathway in Fig. 1c is a closed loop, because the output is fed back into the system.

This section compares the performance costs and benefits of an open loop system without feedback and a closed loop system with feedback.

### Performance metric: plasticity vs homeostasis

To analyze tradeoffs, we need a performance measure. Before choosing a measure, it is useful to consider two attributes with respect to the goals for this section. First, we want a measure that provides sufficient generality to achieve broad insight and also provides sufficient specificity to allow numerical illustration. Second, we want a measure that emphasizes the tradeoff between the responsiveness of plasticity to environmental change and the ability to hold a homeostatic setpoint in response to random perturbations.

Often, a system that responds relatively quickly to environmental change also deviates more easily from a homeostatic setpoint. A tradeoff occurs between responsive plasticity and homeostasis.

Responsiveness can be measured in many different ways. Classic control theory often analyzes how quickly and accurately a system responds to a step change in the environmental reference signal. For example, the environmental signal may initially be a constant value of zero to represent the baseline environment. Then, suppose the environment shifts instantaneously to a constant value of one. How does the system respond to that unit step change?

Using the measure in eqn 10, we can write the step-response performance between an initial time, *t =* 0, and an arbitrary final time, *T,* as

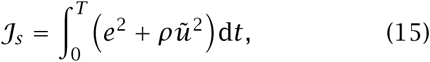

Figure 7a shows how the second-order dynamics in eqn 12 responds to a step change in input. For the various values of the parameter *α* illustrated in the figure, the caption lists the associated performance measure, *𝒥*_*s*_ with *ρ =* 0, which reduces *𝒥*_*s*_ to the integral of the quadratic error, *e*^2^.

**Figure 7:**
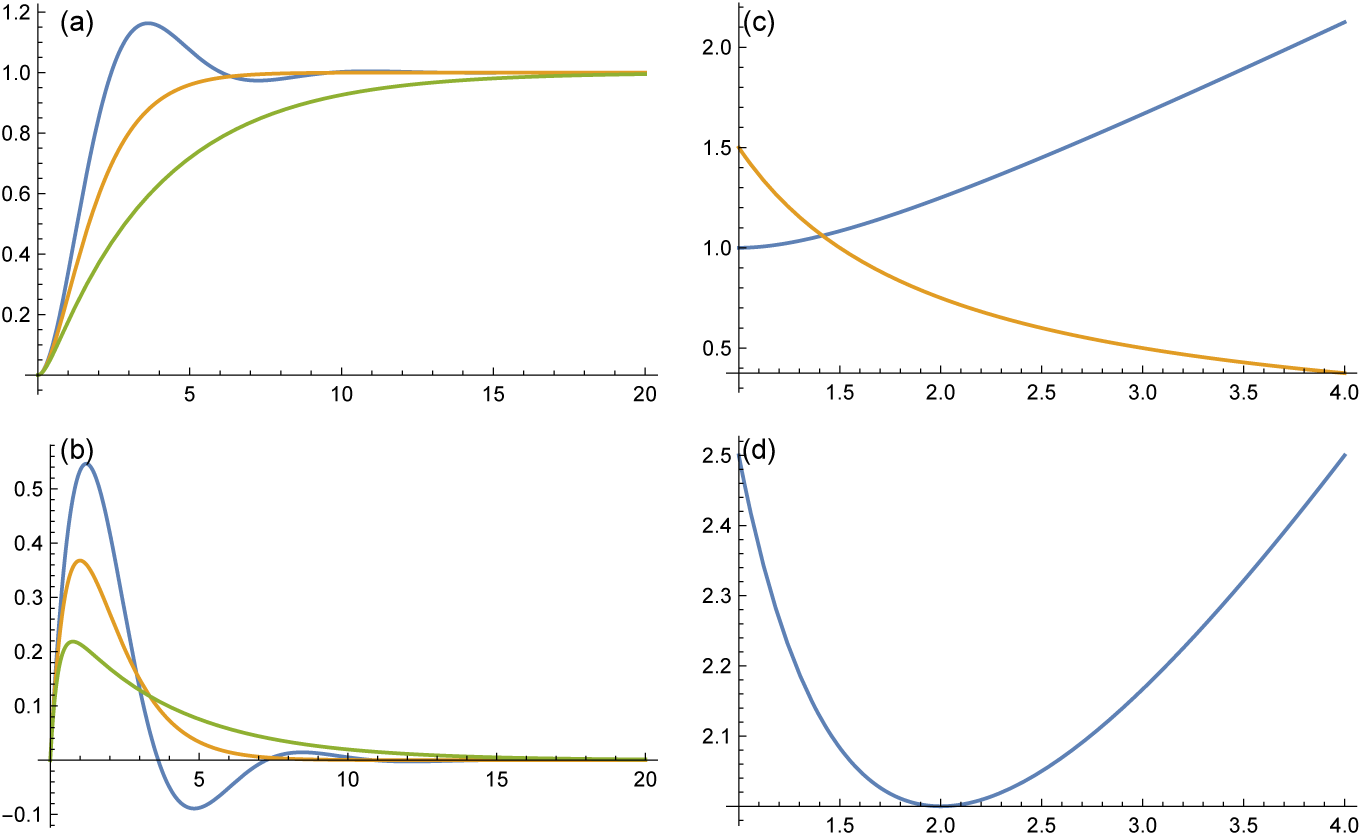
Performance metric for plasticity versus homeostasis. (a) Responsive plasticity to a unit step change in the environmental reference signal. The curves from top to bottom show *α =* 1, 2, 4 from eqn 12. (b) Homeostatic return to setpoint after impulse perturbation. Same underlying dynamics and parameter values as the upper panel. (c) The components of performance given by the cost metrics *𝒥*_*s*_ in blue and *γ 𝒥*_*p*_ in gold for *γ =* 3 and varying values of the parameter *α* along the *x*-axis. Lower values of the cost metrics correspond to better performance. (d) Total performance cost *𝒥 = 𝒥*_*s*_ *+ γJ*_*p*_ for *γ =* 3. The optimal minimum 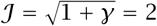 is at 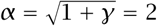.

Homeostasis, or regulation to hold a setpoint, also can be measured in many different ways. Classic control theory often analyzes how a single, large instantaneous perturbation causes a system to deviate from its setpoint, and the path that the system follows as it returns to its setpoint. Technically, at some time instant, say *t =* 0, the system experiences a perturbation to its input of infinite energy and infinitesimal duration, a Dirac delta impulse perturbation. In the typical analysis, the input is a constant value of zero before and after the perturbation. Thus, the error is the output deviation from zero, *y*, and the control deviation from the input is *ũ = ur = u* for input *r =* 0, so we can write the performance as

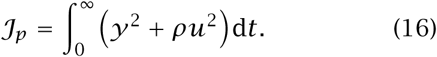

Figure 7b shows how the second-order dynamics in eqn 12 responds to an impulse perturbation in input. For the various values of the parameter *a* illustrated in the figure, the caption lists the associated performance measure, *𝒥* _*p*_ with *ρ =* 0, which reduces *𝒥* _*p*_ to the integral of the quadratic deviation of the output from the zero setpoint.

In signal processing theory, an integral of the squared deviations over an infinite time period, such as 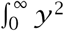 d*t*, is referred to as the energy of a signal. For finite energy signals, an interesting identity provides conceptual and numerical benefits. For the response to an impulse perturbation input, the energy is proportional to a measure, *ℋ*_2_, of the area under the Bode magnitude plot curve, such as in Fig. 2e. Thus, we can use the frequency response of a system to understand and to calculate a system’s intrinsic homeostatic regulation.

The total performance measures the tradeoff between plasticity and homeostasis as

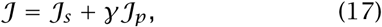

in which *γ* describes the weighting of the homeostatic performance in response to perturbation relative to the plasticity performance in response to a step change in the environmental reference signal. Optimal performance minimizes *𝒥*, as shown in Fig. 7d.

For this example, I weighted the control signal by *ρ =* 0, thus ignoring the cost of control. With that assumption, 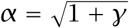 in the second order process *P* in eqn 12 yields the optimal performance value 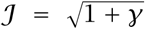, which balances the tradeoff between plasticity and homeostasis. See the supplementary Mathematica code for the derivation of the optimal value of *α* and associated minimization of *𝒥*. In some of the analyses below, I vary *α* around its optimal value as a way in which to introduce environmental perturbations.

### Optimal open and closed loops

This section analyzes the costs and benefits of error feedback. To quantify those costs and benefits, I compare the design and performance of a system without feedback and a system with feedback.

We can denote the system generically from Fig. 1 as *r* → *G* → *y*, in which the *r* is the environmental reference input signal, *y* is the system output, and the transfer function, *G*, represents all of the system components between the input and output signals.

Consider the open loop system in Fig. 1b. We can write that system as *G = CP = L*, in which *C* is the controller transfer function, *P* is the plant transfer function, and *L* denotes the open loop, *CP,* between the input and the output. The closed loop system in Fig. 1c has *G* = *L*/(1 + *L*), as derived in eqn 8. For the plant, I use *P* in eqn 12, which depends on the parameter, *α*.

The analysis proceeds as follows. First, all derivations in this section start by optimizing the performance metric, *𝒥*, in eqn 17, ignoring signal costs, *ρ =* 0.

Second, I use the value of *α* in the plant, *P,* that optimizes the performance metric, *𝒥* in eqn 17, for the uncontrolled system *r* → *P* → *y*. From the previous section, we have the optimal value 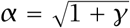 and associated optimal performance 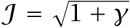. By using the optimal value of *α*, we can consider how a controller improves system performance by altering the constraints on the dynamics rather than by simply tuning the plant’s intrinsic dynamics.

Third, for the controller, I use

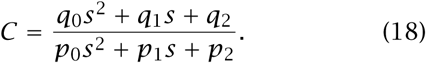

Optimization finds the best values of the *q*_*i*_ and *p*_*i*_ parameters. For the performance metric described in the previous paragraphs, the numerical optimization analysis suggests that, for both open and closed loop systems, the optimal controller typically transforms the uncontrolled second order plant system, *r* → *P* → *y*, into the first order controlled system *r* → *G* → *y*, in which

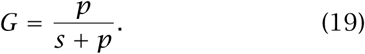

For all numerical optimizations, I used the NMinimize procedure from version 11.3 of Wolfram Mathematica (https://www.wolfram.com/mathematica/) or the differential evolution method from version 2.7 of the Pagmo optimization software library (https://esa.github.io/pagmo2/).

### Optimal controllers

For the open loop system, in which *G = L = CP,* we can write

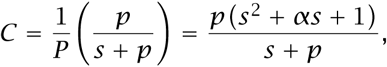

so that in the system *G = CP,* the 1*/P* term in the controller cancels the plant, *P.* In the control optimization of deterministic systems, such canceling of terms often occurs, because the system can reshape response dynamics by removing the fixed plant dynamics and replacing those dynamics with the modifiable dynamics of the controller.

For the closed loop system, in which *L*\**= C*\**P,* we can write

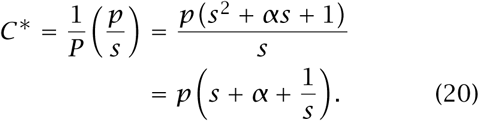

This form is known as a proportional, integral, derivative (PID) controller, the most widely used general controller in systems engineering. From the right-hand side, the term *α* amplifies the input signal by a constant value of proportionality, the term 1*/s* yields the integral of the input signal, and the term *s* yields the derivative of the input signal. These three terms provide wide scope for modulating the dynamics of a feedback system.

The controller *C*\* yields the open loop

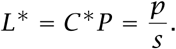

In earlier sections, I discussed how an open loop integrator, *L*\**= p/s*, yields a closed loop low pass filter with exponential dynamics. Thus, we obtain

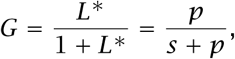

which is a simple first order low pass filter, as in eqn 6. A low pass filter corresponds to basic exponential dynamics, as discussed in earlier sections. These derivations show that the optimizing controllers transform the second order plant, *P,* into the first order low pass filter system, *G*.

The supplementary Mathematica file demonstrates analytically that 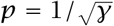 yields the optimal performance 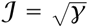. Thus, the optimal controllers shown here improve the optimized plant performance from the uncontrolled value of 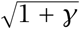 to the controlled value of 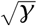. For the analytic analysis, the time period for the step response performance in eqn 15 is *T = ∞*. In numerical analysis, the step response typically converges to the reference input, *r,* by *T =* 20, as illustrated in the various example plots.

Figure 8 compares the dynamics of the unmodified plant, *P,* with the optimized open loop system, *G = CP.* The unmodified plant, shown in the blue curves, has reasonably good response characteristics, because we chose the parameter 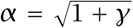 to optimize the performance cost metric. For *γ* = 1, the unmodified plant has performance 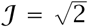. The optimized open loop, shown in the gold curves, improves the response characteristics, yielding an improved performance metric, *𝒥 =* 1.

**Figure 8:**
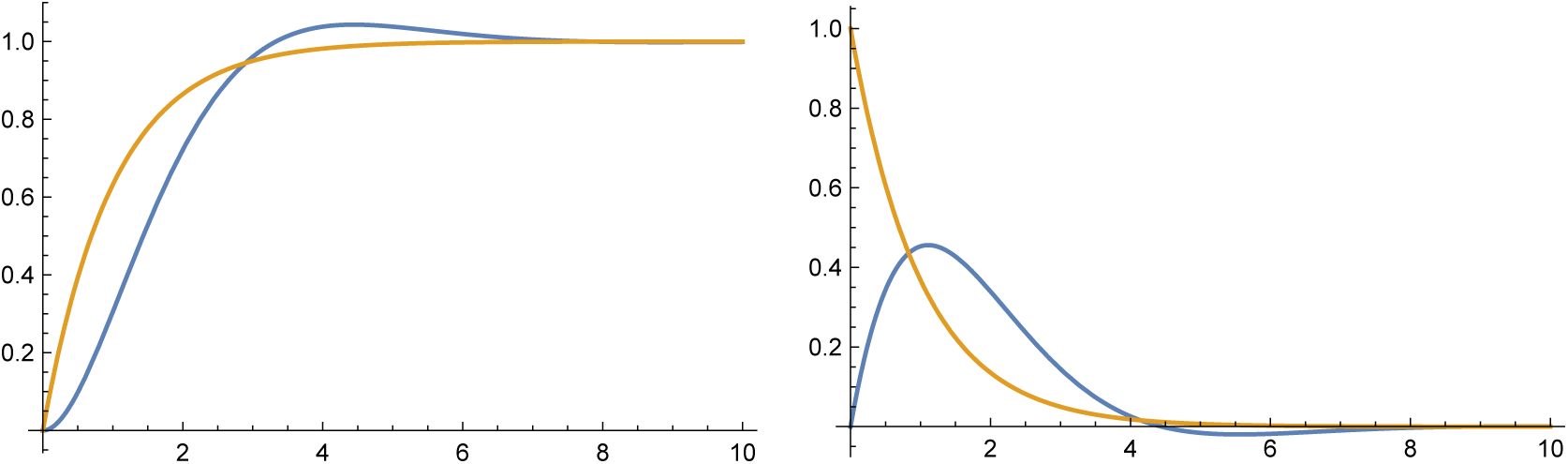
Dynamics of the unmodified plant, *P,* from eqn 12, with *γ =* 1 and optimal parameter 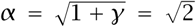 (blue curves) compared with the optimized open and closed loop systems with dynamics given by the low pass filter *G* = *p*/(*s* + *p*) with 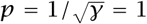 (gold curves). The left panel shows the unit step response, and the right panel shows the impulse perturbation response.

### Comparing open vs closed loop systems

When the plant is a known, deterministic process, the optimal open and closed loop systems yield the same system response. We can write both open and closed loop systems as *r* → *G* → *y*, in which *G* is the open or closed loop processing of the input, *r,* to yield the output, *y*.

Typically, the closed loop system is more costly to build and run, because it requires sending a measure of the output, *y*, back to controller input, and subtracting that output from the reference input, *r,* to produce an error signal, *e = r − y*. The open loop system does not require that extra signal transmission and processing. Thus, for known deterministic systems, open loops generally outperform closed loops.

That general advantage of open loops is a well known principle of engineering control design. As noted by Vinnicombe (2001, p. xvii): “There are two, and only two, reasons for using feedback. The first is to reduce the effect of any unmeasured disturbances acting on the system. The second is to reduce the effect of any uncertainty about systems dynamics.”

Closed loop systems with feedback can often correct for disturbance and uncertainty. With a feedback measure of error, a closed loop system can always do well simply by moving in the direction that reduces the error. By contrast, open loop systems cannot correct themselves because they lack a measure of error. With regard to design principles, the key question concerns how often inevitable uncertainties favor the complexity of closed loop feedback over the simplicity of an open loop design.

### Sensitivity to uncertainty

In this section, I analyze uncertainties in the dynamics of the intrinsic plant process and in the controller. I show that a closed loop design is much less sensitive to those uncertainties than an open loop design.

Figure 9a illustrates how the performance cost metric, *𝒥*, increases as the plant process parameter, *α*, takes on variable values, 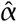, yielding the uncertain plant

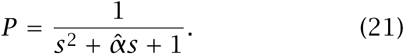

**Figure 9:**
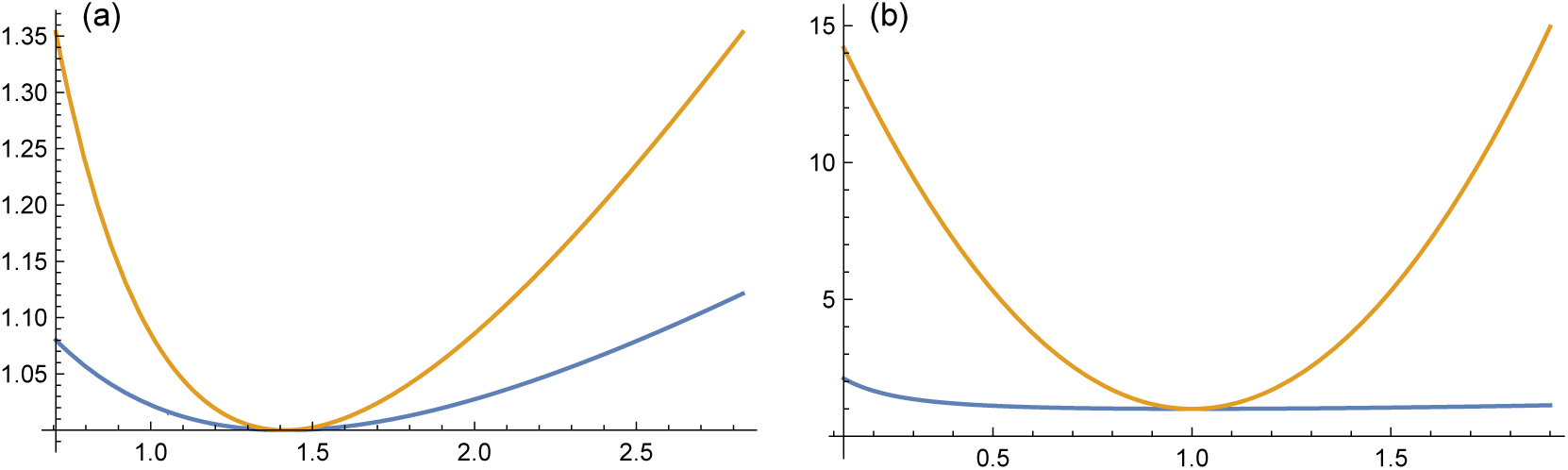
Sensitivity of open loop (gold curve) and closed loop (blue curve) systems to parametric variations in dynamics. The *y*-axis shows the performance cost metric, *𝒥*, and the *x*-axis shows a variable parameter. (a) The plant dynamics parameter, *α*, takes on variable values, 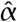. (b) The controller dynamics parameter, *q*_2_, takes on variable values.

The open loop (gold curve) and closed loop (blue curve) take on their identical minimum values at the optimum 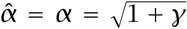, with *γ =* 1 in this example. As 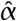 varies, the open loop performs much worse than the closed loop. In other words, the open loop is much more sensitive to variations in plant system dynamics than is the closed loop.

Similarly, Fig. 9b illustrates the greater open loop sensitivity to variations in the controller dynamics. In this case, the controller parameter *q*_2_ takes on variable values around its optimum at 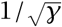.

### General analysis of sensitivity

Closed loop systems are generally less sensitive to parametric variations than are open loop systems. We can study that general pattern of sensitivity by analyzing the derivatives of the systems with respect to parameter variations. In particular, write an open loop system as *L* and a closed loop system as *L*\*/(1 + *L*\*), and let *∂* be the partial derivative with respect to some parameter, *θ*. Then we can write the parametric derivative to describe the relative sensitivity of closed versus open loop systems

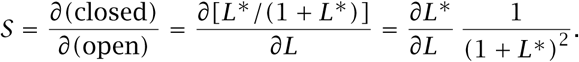

At low frequency inputs, closed loop systems typically have high gain values for their open loop components, *L*\*, causing the closed loop systems to be much less sensitive to parametric variations.

In the example of this section, *L = CP* and *L*\**= C*\**P,* for which *P* depends on the variable parameter 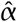, as shown in eqn 21. Taking the partial derivative with respect to 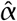, the relative sensitivity of the closed versus open loop systems is

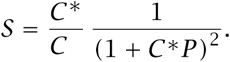

Noting from Fig. 1 that we can express the open loop system as *Y = RCP* and the closed loop system as *Y = REC*\**P,* equating these expressions for *Y* yields *C*\**/C =* 1*/E*. From eqn 9, we have 1*/E =* 1 *+ L*\**=* 1 *+ C*\**P,* thus the relative sensitivity reduces to the simple expression

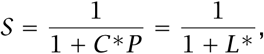

which provides a very general description for the reduced sensitivity of closed loop error feedback systems relative to open loop systems with respect to variations in the plant process, *P.*

For this particular example, we can use the expression for *C*\*in eqn 20 and *P* in eqn 12, yielding

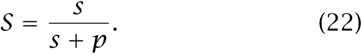

These relative sensitivity expressions further demonstrate the power of transfer functions for the analysis of control dynamics. Using the general properties of transfer functions, we can describe how sensitivity changes with the frequency of inputs to the system. At low frequency, *s* → 0, the relative sensitivity of closed versus open loops declines toward zero, implying a large advantage for closed loop systems versus open loop systems.

In general, we can plot the relative sensitivity on a log scale by using the Bode magnitude plot method in Fig. 2e, yielding the relation between relative sensitivity, *S*, and frequency, as shown in Fig. 10. In that figure, we see a pattern analogous to the classic high pass filter. At high input frequencies, the system has a gain of one (log(1*) =* 0), corresponding to equal sensitivity of open and closed loops. As the input frequency declines, the relative sensitivity of the closed loop declines.

**Figure 10:**
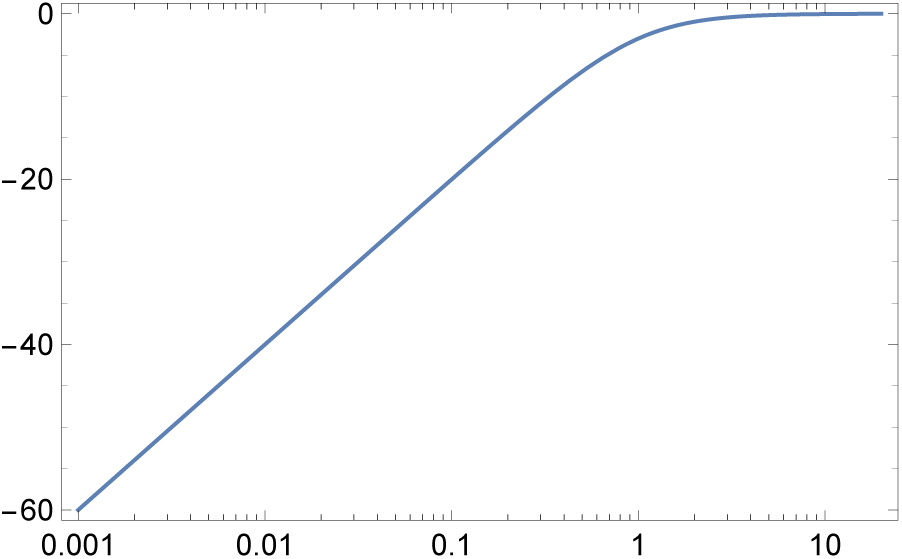
Sensitivity of closed versus open loop systems given in eqn 22. As in Fig. 2e, this Bode plot shows the logarithm of the transfer function magnitude versus the logarithm of the input frequency.

### Frequency tradeoffs

The sensitivity pattern in relation to frequency illustrates a key point about control system design. When considering tradeoffs, we must often analyze how tradeoffs change in relation to the frequency of inputs and the frequency of perturbations to a system. Often, improving performance at one frequency band reduces performance at another frequency band, creating an additional perspective on the tradeoffs in design.

The essence of the plasticity versus homeostasis tradeoff comes down to a tradeoff between an organism’s response to different frequencies of inputs. In the examples here, plasticity associates with the response to a step change in input, and homeostasis corresponds to the response to an impulse perturbation.

Consider the response to a step input in terms of frequency. A step input has a transfer function 1*/s*, in which the input has a lot of energy at low frequency (small *s*) and decreasing energy as frequency increases. If an organism filters out high frequency inputs and responds to low frequency inputs, it will slowly and accurately adjust to long-term environmental change. The more strongly the organism responds to high frequency inputs, the more rapidly the organism adjusts to sudden step changes in environmental state. We may say that greater high frequency sensitivity corresponds to a more malleable and rapid plastic response.

The plasticity versus homeostasis tradeoff arises because the greater the plastic response to the high frequency components of step inputs, the more strongly the organism’s homeostasis will be disrupted by impulse perturbations. In particular, an impulse input has a constant transfer function, which corresponds to an input with equal energy over all frequencies. The wider the frequency bands that the organism filters out and does not respond to, the less sensitive the organism is to impulse perturbations that disturb homeostasis.

Putting all of that together, responding to low frequencies is most important for plasticity, because step inputs with transfer function 1*/s* strongly weight the lower frequencies. By contrast, homeostasis gains by filtering all frequencies equally, because impulse energy is uniformly distributed over all frequencies. Thus, to balance good plasticity and good homeostasis, the organism must respond to inputs in the lower frequency band and reject inputs in the higher frequency band.

As *γ* declines, the performance metric weights the plastic step response more strongly than the homeostatic perturbation response. Thus, a decline in *γ* favors the organism to increase its range of high frequency sensitivity, to achieve the benefit of faster, more malleable plastic response while paying the relatively low cost of suffering greater homeostatic perturbation.

The parameter 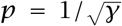 in eqn 19 corresponds to the point at which the system more strongly filters relatively higher frequencies. As *γ* declines and performance weights plasticity more strongly than homeostasis, the optimal organismal design moves the frequency cutoff *p* to higher values, causing the organism to respond to relatively higher frequency inputs.

### Sensitivity and variability

The closed loop system is less sensitive to parameter variations than the open loop system. Less sensitivity means that mutations have less effect on performance. Thus, the theory predicts that the closed loop systems will accumulate more genetic variability than the open loop systems (Frank, 2004, 2007).

Less sensitivity also means that a system can tolerate greater stochasticity in phenotypic expression with less consequence for performance. Thus, the theory predicts that the closed loop systems will have greater phenotypic stochasticity in gene expression than the open loop systems (Frank, 2013).

A subsequent article in this series develops the prediction that low sensitivity and high robustness enhance genetic and phenotypic variability.

### Summary

Error feedback compensates for uncertainty and stochastic perturbation. Because most biological systems have uncertain or stochastic aspects, feedback control inevitably plays a key role in biological design. However, the net benefit of feedback depends on the frequency of system inputs and on the various costs associated with running the relatively complex control loop of error correction.

One aspect of cost arises from the extra components and energy required to build and run the feedback loop. Another aspect of cost concerns the strength of the control signal required to correct errors. The next section considers the costs associated with control signal strength.

## Costs of control

In the prior section on optimizing open and closed loops, I ignored the cost of the control signal. This section extends that prior optimization to include the control costs, by assuming that the control cost weighting is greater than zero, *ρ >* 0.

The general performance cost metric from eqn 17 is *𝒥 = 𝒥*_*s*_ *+ γ 𝒥*_*p*_. The step input response component, *𝒥* _*s*_, was defined in eqn 15 as

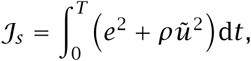

for which *ũ = u-r* is the difference between the control signal, *u*, and the environmental reference input signal, *r.* For a step input, *r =* 1. The control cost depends on how much the control process causes a deviation from the external environmental state.

Similarly, the impulse perturbation response component *𝒥* _*p*_ was defined in eqn 16 as

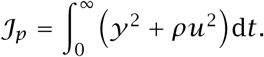

For an impulse response, the environmental reference signal is zero except for an infinite impulse over an infinitesimal duration around time *t =* 0, so *ũ = u r ≡ u*. In the numerical calculations of the control signal cost, I ignore the infinitesimal impulse interval, and sum up the deviations over the period *t >* 0, after the impulse.

### Control signal dynamics

To analyze the control costs, we need expressions for the dynamics of the control signal, *u*. For the open loop system in Fig. 1b, the control signal is simply the response of the controller to the reference input. Thus, *U = C*, and we can study the control signal response by analyzing the controller transfer function.

For the closed loop in Fig. 1c, the control signal arises as the response of the controller, *C*, to the error input signal, *E*, thus *U = EC*. From eqn 9, *E =* 1*/(*1 *+ L)*, in which *L = CP.* Thus the control signal is *U* = *C*/(1 + *L*).

In deterministic systems with known dynamics, the optimal open and closed loop systems will produce the same control signal. Thus, the optimal controller for the open loop system, *C*, will be related to the optimal controller for the closed loop system, *C*\*, by the relation *C = EC*\*.

The error component, *E*, typically reduces the magnitude of the reference input, *R*. Thus, the optimal closed loop controller by itself tends to amplify signals much more strongly than the optimal open loop controller.

In terms of biology, this theory predicts that controlling subsystems designed to modulate internal biochemical signals within a broader system will amplify signals in very different ways when in open versus closed loops.

### Zero cost pass-through controller

Before studying the role of variable costs in shaping the tradeoff between plasticity and homeostasis, it is useful to consider the special case of high control signal costs.

If the cost weighting, *ρ*, for the control signal is very high, then the optimal system will reduce the control signal deviation *ũ = u r* to zero, which means *u = r.* In that case, the optimal controller will simply pass through the unchanged reference signal, and the optimized control signal is *U =* 1. In this case, the open loop controller is simply *C =* 1.

For closed loops, noting that *L*\**= C*\**P,* the closed loop controller is

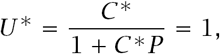

thus

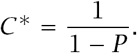

It seems likely that as *ρ* increases, the benefit of the closed loop system with regard to plant uncertainty and disturbance rejection declines until that benefit disappears at the pass through limit for large *ρ*. The following section analyzes that conjecture.

## Plasticity vs homeostasis

We can now consider quantitative examples of two key tradeoffs in control system design. First, systems typically must trade off plastic responsiveness to changing environments versus homeostatic buffering against perturbations. Second, systems trade off the costs of producing control signals to adjust dynamics versus the benefits of those control signals to manipulate dynamics.

### Performance metric expression of tradeoffs

The prior sections developed a performance metric that captures the two tradeoffs. The full expression of the performance metric can be written as

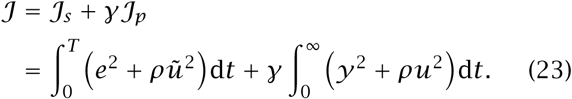

The first line emphasizes the plasticity versus homeostasis tradeoff. The parameter *γ* weights the relative importance of plastic responsiveness, given by the step response performance, *𝒥*_*s*_, and homeostatic buffering, given by the perturbation response, 𝒥_*p*_. Smaller values correspond to higher performance. Thus, a greater value of *γ* favors improving homeostatic performance at the expense of reduced plasticity.

The second line emphasizes the tradeoff associated with control signal strength. A greater value of *ρ* favors reducing the control signal strength. Reduced control signal strength may alter the tradeoff between plasticity and homeostasis, as illustrated in the following example.

### Numerical example

This example begins with the optimized plant in eqn 21, with 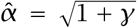. The general form of the controller is given in eqn 18. For each combination of *γ* and *ρ* parameters, I used the differential evolution method of Pagmo 2.7 to find a candidate combination of controller parameters that optimizes (minimizes) the performance metric in eqn 23.

The optimal controllers differ for open and closed loops. However, when the system is deterministic and there is no uncertainty in the plant dynamics, the optimized open and closed loops have identical dynamics and control signals, as described in the prior section. Thus, this example does not differentiate between optimized open and closed loops. The following section develops this example when the plant parameter *α* varies, which highlights the differences between open and closed loops under uncertainty.

We can partition the total performance metric in eqn 23 into four components. From *𝒥*_*s*_, the components in the step response are: (a) the squared error deviation from the environmental setpoint, *e*^2^, and (b) the squared control signal deviation from the signal that would occur in a pass-through controller, *ũ*^2^. From *𝒥*_*p*_, the components in the perturbation response are: (c) the squared deviation from the base-line zero setpoint, *y*^2^, and (d) the squared control signal deviation from zero, *u*^2^. The various components are weighted by the parameters *γ* and *ρ*.

Figure 11 shows the components of performance in optimized systems. Lower values correspond to better performance. Panels (a–d) present the four components of performance in the order given in the prior paragraph. Each panel plots log_2_ of the component performance versus log_2_ *γ*. The curves in each panel show log_10_ *ρ =* −4, −3,*…,* 0. In panel (a), the values of *ρ* rise from the bottom to the top curve, whereas in the other panels, the values of *ρ* decline from the top to the bottom curve.

**Figure 11:**
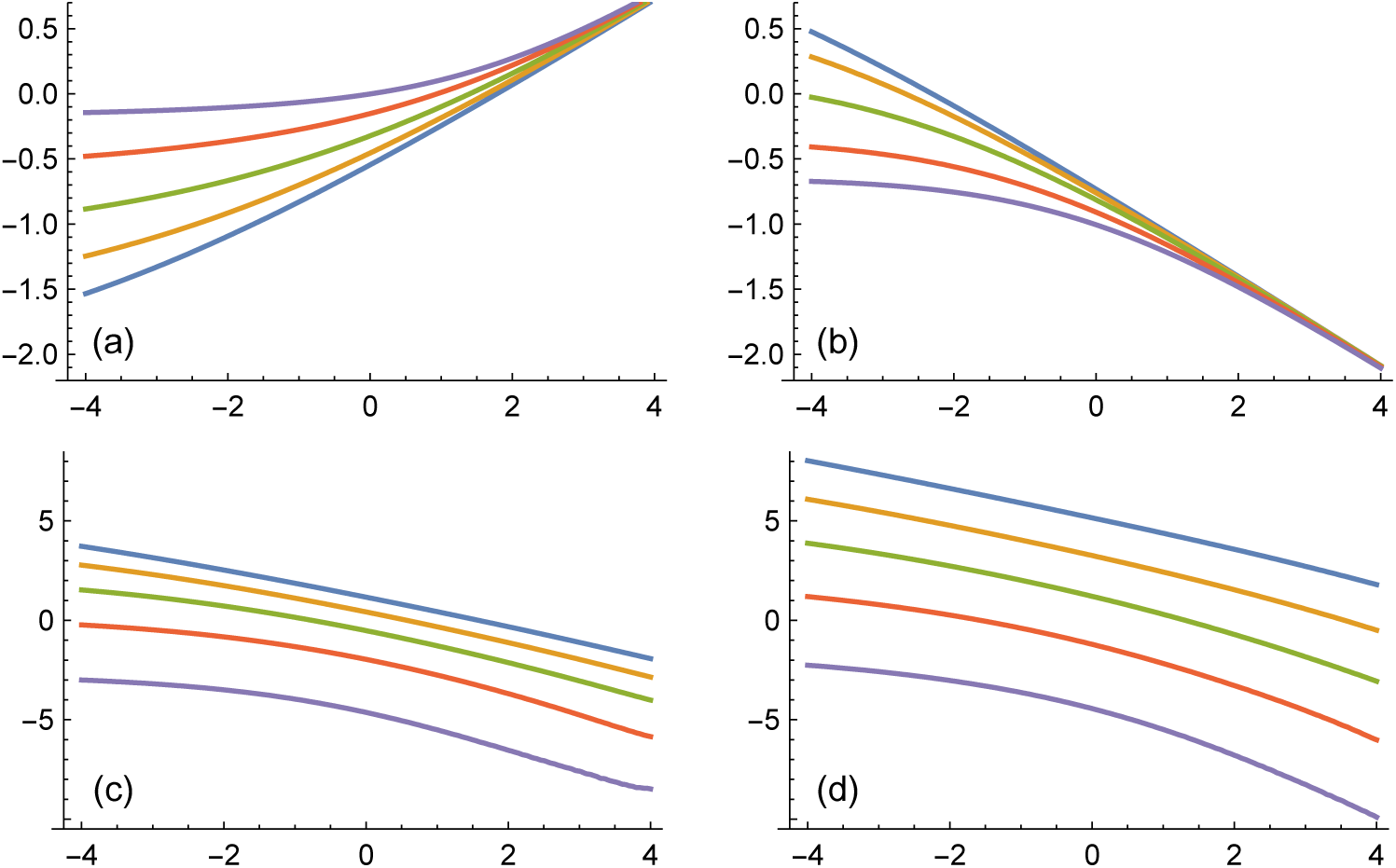
The tradeoffs between plasticity, homeostasis, and the costs of control signals. Each panel shows log_2_ of a component of performance versus log_2_ *γ*. The text describes the different components of performance. Within each panel, the five curves correspond to log_10_ *ρ =* −4, −3,*…,* 0. In panel (a), the values of *ρ* rise from the bottom to the top curve, whereas in the other panels, the values of *ρ* decline from the top to the bottom curve.

In Fig. 11, comparing panels (a) and (b) illustrates the primary plasticity versus homeostasis tradeoff. As *γ* increases, the performance metric depends more strongly on homeostasis, causing the homeostasis component to show improved (lower) contributions to *𝒥* and an associated rise in the plasticity component. In essence, the system responds more slowly to environmental changes, which benefits homeostatic performance with respect to perturbations and weakens plastic responsiveness to a rapid, long-term shift in the environment.

As the cost of the control signal rises, the system shifts toward improved homeostatic performance at the expense of reduced plastic responsiveness. In the homeostatic response of panel (b), rising signal cost, *ρ*, corresponds to lower (better) performance curves. In the plastic response of panel (a), the opposite pattern of higher (worse) performance associates with rising signal cost.

The plasticity versus homeostasis tradeoff with rising signal costs occurs because high signal costs tend to favor a lower control signal intensity and thus a slower system response. A slower system reduces the rate of plastic responsiveness to a sudden, longterm environmental shift and also reduces the sensitivity of the system to short-term perturbations that disturb homeostasis.

Panels (c) and (d) show the inevitable decline in control signal energy with rising signal cost. The higher signal energies in panel (d) reflect the strong system response to the high-intensity instantaneous jolt to the system caused by the perturbation.

## Open vs closed loops

The previous section assumed that the parameters controlling the intrinsic plant dynamics are known and constant. For known plant dynamics, optimized open and closed loops perform identically. If plant dynamics deviate from the assumed form, open and closed loops typically perform differently.

This section considers the relative performance of open versus closed loops for variable plant dynamics. As in previous sections, I describe variable plant dynamics with eqn 21, in which the optimized plant has parameter 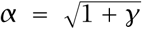. The open and closed loops are then optimized with respect to the optimized plant. I then analyze the open and closed loop responses when the plant parameter 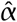 varies around the optimum *α*.

Figure 12 illustrates the relative performance of open versus closed loops as 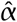 varies over (*α*/2, 2*α*). The curve plots log_2_ *𝒥*_*o*_*/ 𝒥*_*c*_, the log ratio of the open loop performance relative to the closed loop performance. The curve touches zero at 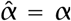. For other values of 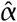, the closed loop outperforms the open loop, which means that the closed loop has a lower *𝒥* value than the open loop.

**Figure 12:**
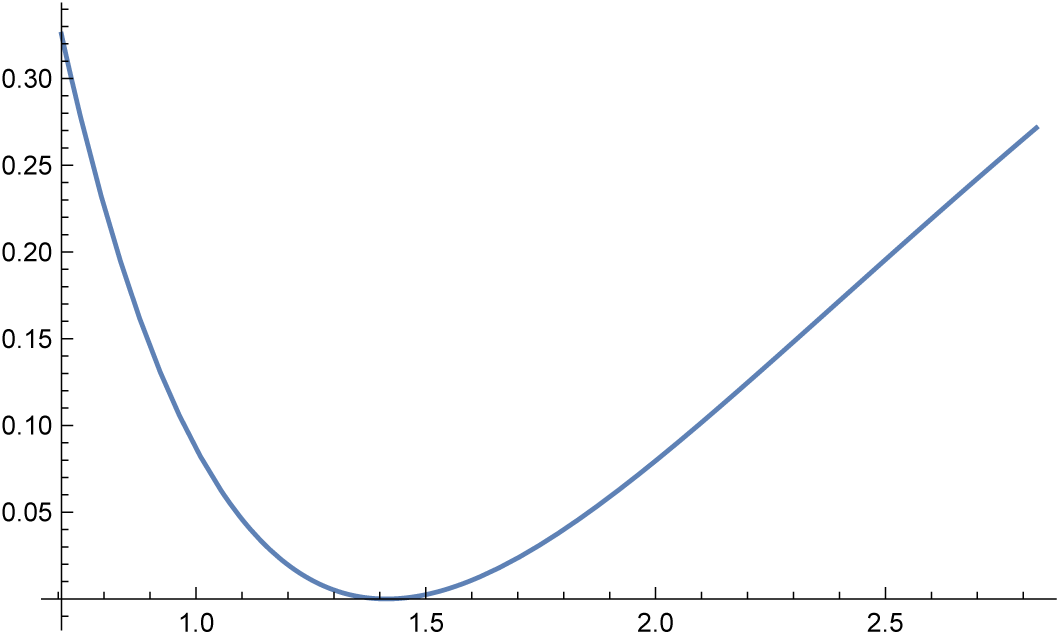
Log ratio of open versus closed loop performance for variable plant dynamics. The *x*-axis shows the varying plant parameter 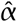 from eqn 21.

Figure 12 shows the same information as Fig. 9a, but plotted in a different way. The advantage of Fig. 12 is that we can take the average performance ratio over the range of 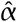 values as a rough measure of the relative advantage of the closed loop versus the open loop for variable plant dynamics. The averaged relative performance metric is the integral area under the curve in Fig. 12 divided by the range of 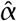 values. For Fig. 12, the average height of the curve is 0.108.

Plotting Fig. 12 requires specific parameter values for *γ* and *ρ*, which determine the relative weighting of various performance components. In that figure, *γ =* 1 and *ρ =* 0.

Figure 13 plots the averaged performance metric for varying values of *γ* and *ρ*. The *x*-axis shows log_2_ *γ*. The curves show varying values of *ρ*. The top curve is for *ρ =* 0. The subsequent curves are for log_10_ *ρ =* −4, −3, −2, −1.

**Figure 13:**
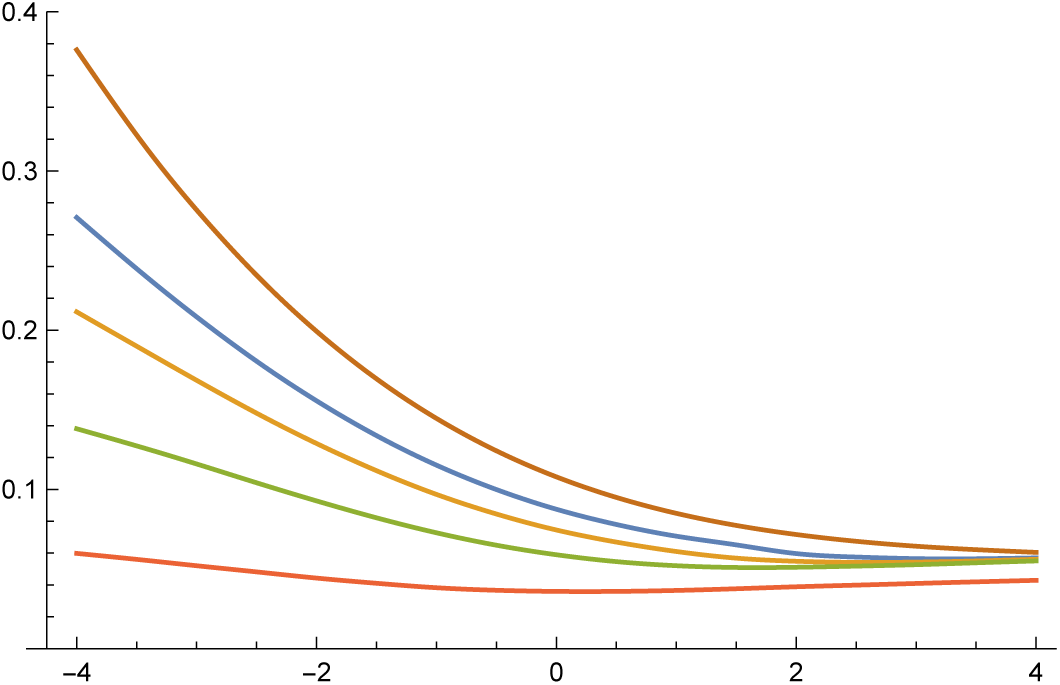
Log ratio of open versus closed loop performance when averaged over variable plant dynamics. The *x*-axis shows 2^*γ.*^ The curves from top to bottom plot results for increasing values of *ρ*.

Closed loops perform relatively better for small *γ*, corresponding to greater emphasis on plastic responsiveness to sudden onset, long-term changes in the environment. Because closed loops use the error as input, they strongly drive the system toward a new setpoint when far from that setpoint and smoothly ease up on the push toward the setpoint as the gap is closed. Closed loops also perform better for lower values of *ρ*, because they benefit from strong control signals to drive the system when far from the target setpoint.

For the parameters considered here, closed loops outperform open loops. However, closed loops require additional components, internal measurements, and signal transduction, which may add additional costs. With those additional costs, simpler open loop systems will tend to outperform closed loops when there is relatively little uncertainty, when control signals are costly, and when homeostatic rejection of perturbation is more heavily weighted than plastic responsiveness (high *γ*).

## Performance vs stability under uncertainty

Instability often causes a system to fail. To protect against instability, a well designed system may trade lower performance in return for broader stability against perturbations or uncertainties.

For example, previous sections optimized a controller solely for performance with respect to the fixed dynamics of a given plant. In this section, I consider optimized controllers subject to the constraint that they must remain stable to a broad range of alternative plant dynamics. A broader stability margin reduces performance of the system for the target dynamics of the given plant.

In the previous analyses, I began with the plant

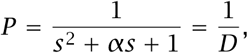

with 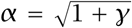 and *D = s*^2^ *+ αs +* 1. I then found optimal controllers with respect to the fixed dynam ics of this plant. This section extends that analysis by requiring that the optimized system also be stable with respect to the alternative plant, 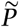, with 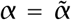, for a given value 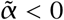.

A system is stable if the real part of its maximum eigenvalue is less than zero. The eigenvalues of a system described by its transfer function are the roots of the characteristic polynomial in *s* in the denominator of the transfer function. The alternative plant, 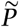, is unstable for 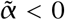 because, by the Routh-Hurwitz stability criterion, all coefficients of a quadratic must have the same sign to be stable.

An open loop essentially cannot stabilize an unstable plant, whereas a closed loop can sometimes stabilize an unstable plant. In particular, suppose we have a controller *C = n/d*, in which the numerator *n* and the denominator *d* are polynomials of *s*, and *d* is stable with maximum real eigenvalue less than zero. The open loop for the alternative plant is

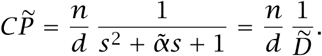

The roots of the denominator will include the roots of 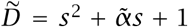, thus the system is unstable unless we can get rid of this quadratic component from the alternative plant. Suppose that 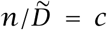 for a constant, *c*, in other words, that *n* cancels 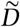. That cancellation removes the eigenvalues associated with *D*, and the system is now stable because *d* is stable.

Although such eigenvalue (pole) cancellation works in theory, it is difficult in practice to achieve sufficiently accurate cancellation, particularly in biological systems. In essence, an unstable plant cannot be stabilized by an open loop controller. Because open loop systems cannot satisfy the constraint of stabilizing alternative plants with 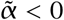, we consider open loops to fail for this constrained optimization problem.

For a closed loop, the system with the alternative plant is

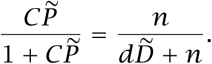

This system is stable if we can find a controller with numerator, *n*, and denominator, *d*, that stabilizes the characteristic polynomial, 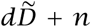. For a controller such as in eqn 18, we will often be able to find stabilizing parameter combinations.

For the closed loop, the optimization problem is to find the best performing controlling for the system with the target plant, *P,* subject to the constraint that that controller also stabilizes the alternative plant, 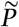 with parameter 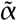.

As a first step, consider a closed loop optimized for performance with respect to the target plant, *P,* without any stability constraint for the alternative plant, 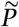. For this case, Fig. 14 shows the value of 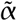 at which the optimized closed loop transitions to instability, with systems below the line being unsta ble.

**Figure 14:**
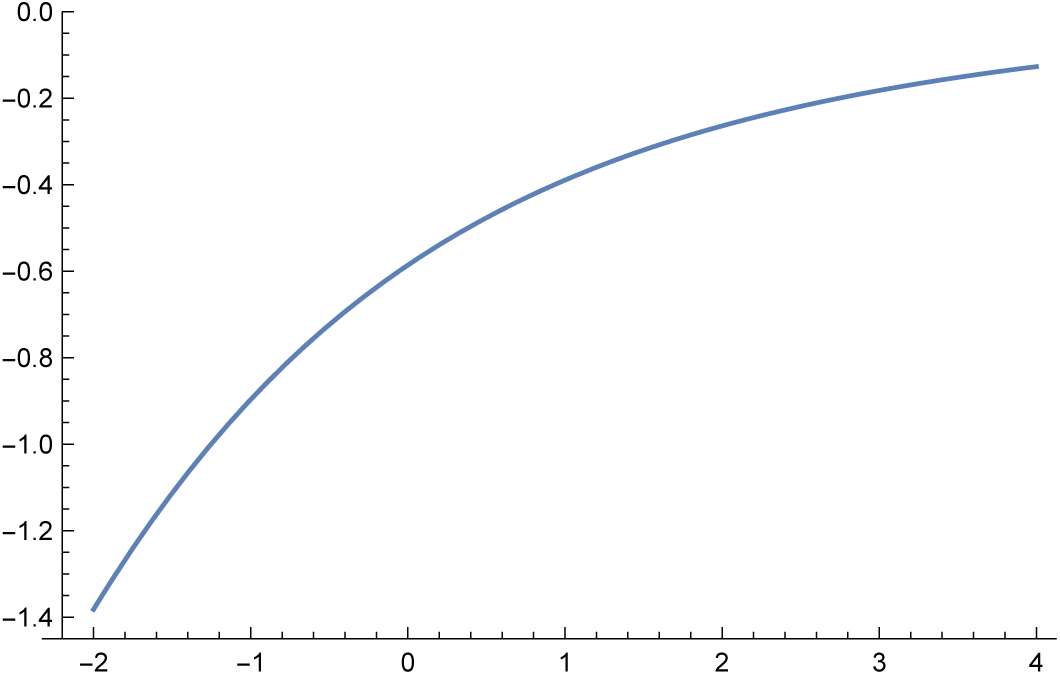
Value of 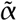 in an alternative plant, 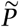, that destabilizes a closed loop optimized for the target plant, *P.* The *x*-axis shows the value of log_2_ *γ* used in the optimized performance metric.

Note that the transition depends on *γ*, which weights the relative importance in the performance metric of plastic responsiveness to a changed environment versus homeostatic performance with respect to sudden perturbations. Larger *γ* weights homeostatic performance more heavily, which favors a slower system response. In this case, slower systems are apparently not as good at stabilizing alternative plants as are faster systems or, put another way, the stronger responsiveness of faster systems apparently stabilizes systems more successfully than slower systems.

For 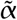 values below the curve in Fig. 14, the unconstrained optimization of closed loops creates unstable systems with respect to the alternative challenge. The next step optimizes the closed loop system for the target plant, *P,* subject to the constraint that the system must be stable with respect to the alternative plant, 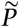, with value 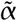.

For 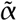 values below the curve, the constrained optimization will reduce the performance of the closed loop with respect to the target plant, *P.* The reduced performance reflects the tradeoff between the performance in a target environment and the safety margin of stability with respect to alternative environments.

Figure 15 illustrates the reduced closed loop performance to achieve a given margin of stability. Each curve shows log_2_ of the ratio between the performance of a closed loop optimized subject to the constraint that it must be stable for a particular 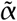 value relative to a closed loop optimized without a stability margin constraint. The curves from bottom to top are for 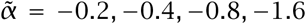. The *x*-axis measures log_2_ *γ*.

**Figure 15:**
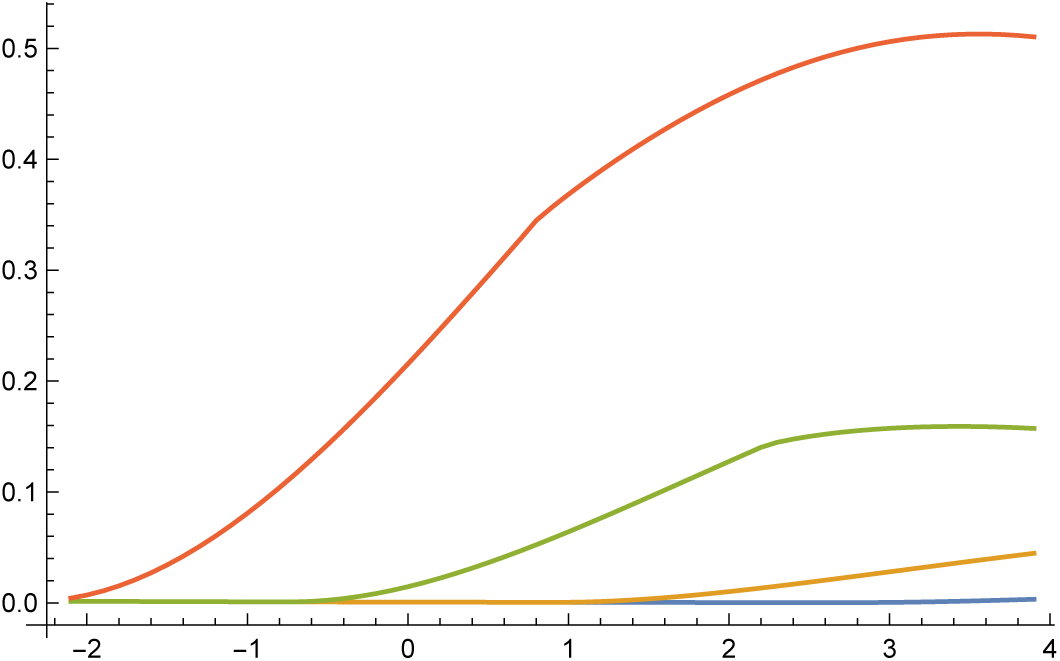
Relative performance of a closed loop optimized subject to a stability margin constraint compared with a closed loop optimized without a stability margin constraint. The plot shows log_2_ of the performance ratio. The curves from bottom to top show an increasing stability margin constraint, corresponding to 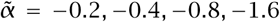. The *x*-axis is log_2_ *γ*.

As the required stability margin becomes greater with declining 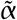 values, the performance cost increases to achieve the required stability margin. As *γ* rises, performance depends more strongly on homeostatic response to perturbation relative to plastic response to a changed environment. The homeostatic response imposes a much stronger tradeoff between stability and performance than does a plastic response.

In this case, a faster system response improves the plastic response and the stability margin but degrades the homeostatic performance. A fast system suffers a greater deviation from the homeostatic setpoint in response to perturbation than does a slow system. In other cases, a faster system response may cause overshoot of the target setpoint, reducing stability.

This section used system eigenvalues to develop the tradeoff between performance and stability margin. Engineering control theory includes more sophisticated methods for the study of stability margins (Vinnicombe, 2001; Frank, 2018a).

## Conclusions

This article introduced the conceptual and analytic foundation for the evolutionary design of regulatory control. The theory provides broad, abstract predictions about various tradeoffs. Those tradeoffs include the balance among the plastic responsiveness of environmental tracking, the homeostatic rejection of perturbations, system stability, and the costs of controls that modulate dynamics. Additional tradeoffs arise from robustness to unpredictable challenges and from alternative responses to different frequencies of inputs.

This broad framework provides the tools to make sense of disparate studies and to develop novel predictions about regulatory control. To give one example of an interesting prediction, consider the general consequences of error-correcting feedback.

Error correction within a system compensates for fluctuations in the performance of the system’s components. That intrinsic robustness of feedback weakens the direct selective pressure on individual components of a system. Weakened selective pressure on components likely increases their genetic variability and their stochasticity of expression.

Although I have discussed those ideas in prior publications, there has been limited work on how control architecture influences the selective pressure on components and the broad consequences for biological variability (Frank, 2004, 2007, 2013). The second article in this series builds on the framework developed here to analyze genetic variability and stochasticity of expression in relation to alternative control architectures (Frank, 2018b).

Another interesting problem concerns the differences between the control architecture of human-engineered systems and the regulatory networks within genomes (Frank, 2017). Eukaryotic gene expression is influenced by transcription factors, methylation, histone codes, DNA folding, intron sequences, RNA splicing, noncoding RNA, and other factors. Vast wiring connectivity links genomic influence to a trait.

An engineer following classic principles of control theory would design a simpler system with fewer connections. Genomes are overwired. They have far more nodes and connections than classically engineered systems.

Why are genomes overwired? It helps to consider what other sorts of systems are overwired. Computational neural networks in artificial intelligence stand out. Deeply, densely connected computational networks pervade modern life. New computational systems often outperform humans.

The recent computational concepts and methods comprise deep learning (Goodfellow et al., 2016). The learning simply means using data, or past experience, to improve the classification of inputs and the adjustment of response. The deep qualifier refers to the multiple layers of deep and dense network connections. That wiring depth, and the computational techniques to use vast connectivity, triggered revolutionary advances in performance.

How can we understand the differences between classic control architecture and the actual wiring of genomes and computational neural networks? The right approach remains an open problem. It seems likely that the greater wiring complexity of genomes and computational neural networks ultimately reduce to fundamental principles of classical control, but also depend on various additional aspects.

One possibility is that the trial and error inductive building of control systems by evolutionary dynamics or computational learning gain from diffusely and densely wired networks, ultimately achieving similar function to classic control architectures but constructed by a different route. Further study along these lines will be interesting.

## Acknowledgments

National Science Foundation grant DEB–1251035 and the Donald Bren Foundation support my research. I completed this work while on sabbatical in the Theoretical Biology group of the Institute for Integrative Biology at ETH Zürich.

## Supplemental files

The Mathematica and Pagmo C++ code are available at: https://doi.org/10.5281/zenodo.1254628.

